# 12-oxophytodienoic acid reductase 3 (OPR3) functions as NADPH-dependent α,β-ketoalkene reductase in detoxification and monodehydroascorbate reductase in redox homeostasis

**DOI:** 10.1101/820381

**Authors:** Daniel Maynard, Vijay Kumar, Jens Sproß, Karl-Josef Dietz

## Abstract

Arabidopsis (*Arabidopsis thaliana*) 12-oxophytodienoic acid reductase isoform 3 (OPR3) is involved in the synthesis of jasmonic acid by reducing the α,β-unsaturated double bond of the cyclopentenone moiety in 12-oxo-phytodienoic acid. Recent research revealed that jasmonic acid synthesis is not strictly dependent on the peroxisomal OPR3. In addition, OPR3 is able to reduce trinitrotoluene suggesting that the old yellow enzyme homologue OPR3 has additional functions. Here we demonstrate that OPR3 catalyzes the reduction of a wide spectrum of electrophilic species that share a reactivity towards the major redox buffers glutathione (GSH) and ascorbate (ASC). Furthermore, we demonstrate that OPDA reacts with ascorbate to form an ASC-OPDA adduct, but in addition OPR3 has the ability to regenerate ASC from monodehydroascorbate (MDHA). The presented data characterize OPR3 as a bifunctional enzyme with NADPH-dependent α,β-ketoalkene double bond reductase and monodehydroascorbate reductase activities (MDHAR). *opr3* mutants exhibited a slightly less reduced ASC pool in leaves in line with the MDHAR activity of OPR3 *in vitro*. These functions link redox homeostasis as mediated by ASC and GSH with OPR3 activity and metabolism of reactive electrophilic species (RES).

## Introduction

12-Oxophytodienoic acid reductases (OPR) belong to the old yellow enzyme (OYE) family and utilize NADPH and flavin mononucleotide (FMN) for reduction of the double bond of the jasmonic acid (JA) precursors 12-oxophytodienoic acid (12-OPDA) and 4, 5-didehydro-JA (ddh-JA) (Vick and Zimmerman 1986; Schaller and Weiler 1997; Chini et al. 2018). Both belong to the group of reactive electrophilic species (RES) and are oxylipins. 12-OPDA and ddh-JA are synthesized from ω-3, 6, 9 polyunsaturated fatty acids (PUFAs) by 13-lipoxygenase (LOX)-dependent oxidation. Subsequently they are modified by additional chloroplastic and peroxisomal enzymes. The products of the oxylipin pathway (especially 12-OPDA and JA) and OPR3 are crucial players in the wound response, flower development, stress acclimation and many other developmental processes in *planta* (Wasternack and Feussner, 2017). Further, oxylipin signaling was linked to antioxidant defense since both 12-OPDA and JA stimulate expression of antioxidant genes under different stress conditions (Maynard et al. 2018b). In particular, several of the identified 12-OPDA- (ORGs) and JA-regulated genes (JRGs) from Arabidopsis (*Arabidopsis thaliana*) are involved in synthesis of glutathione (GSH) and ascorbate (ASC) which are major redox buffers of the cell (Sasaki-Sekimoto et al. 2005; Taki et al. 2005). Apparently OPR3 interferes with redox metabolism through jasmonates.

GSH and ASC play crucial roles in maintaining cellular redox balance (Foyer and Noctor, 2011). Therefore, not only their cellular biosynthesis but also the redox states of the GSH/GSSG and ASC/DHA or MDHA redox pairs are highly regulated through activities of a set of enzymes like glutathione reductase (GR; 1.8.1.7), glutathione-S-transferases (GST; 2.5.1.18), ascorbate peroxidase (APX; 1.11.1.11), dehydroascorbate reductase (DHAR; 1.8.5.1) and monodehydroascorbate reductase (MDHAR; 1.6.5.4) (Foyer and Noctor 2011). Besides their importance as direct or indirect scavengers of reactive oxygen species (ROS), GSH and/or ASC also act as Michael donors and hence react with cytotoxic α,ß-unsaturated carbonyl RES-compounds (Kesinger and Stevens 2009). These Michael reactions where conjugation of a nucleophilic agent (Michael donor) with an electrophilic compound (Michael acceptor) occurs are common and can have protective function in stressed plants. These electrophilic species (RES) are normally generated by processes like non-enzymatic (e.g., phytoprostanes) or enzymatic (e.g., OPDA) oxygenation of fatty acids, secondary metabolic pathways, or in haem metabolism reactions (Farmer and Davoine 2007). RES molecules besides being considered important as stress signals in initial stages of their synthesis are potentially hazardous to cellular constituents due to their high reactivity with nucleophilic atoms (S, N etc.) in proteins or nucleic acids. Therefore, the understanding of interaction of RES and RES-degrading proteins with redox-buffers is of physiological relevance (Almeras et al. 2003; Mueller and Berger 2009).

Although OPR3 is known to reduce the oxylipin and RES 12-OPDA, its function in RES/ROS detoxification has not been addressed in detail. Since JA-synthesis does not show a strict dependency on OPR3 and, in addition, OPR3 possesses the capacity to detoxify molecules like trinitrotoluene (TNT) (Chini et al. 2018) it was proposed that OPR3 has alternative functions that still need to be identified (Beynon et al. 2009). OPR3 protein and activity has been localized to peroxisomes (PaxDB top ten proteins; pax-db.org, TAIR).

Peroxisomes contain high concentrations of ASC and GSH in the millimolar range (Jimenez et al. 1997; Zechman 2011; Corpas and Barroso 2018). The recently described binding of 12-OPDA to GSH (Davoine et al. 2006; Dueckershoff et al. 2008) and the identification of an OPDA-ASC adduct in this study led us to hypothesize and test a role of OPR3 in both RES- and ROS-management. Such a function could be of importance in enzymatic RES-detoxification (Palma et al. 1991; Fryer 1992; Farmer et al. 2003; Kao et al. 2018) besides reducing oxidative damage under stress conditions. In this context the study addresses potentially new functions of OPR3 and provides novel insight into biochemical activities of OPR3. It appeared remarkable that OPR3 can regenerate ASC from monodehydroascorbate, besides having broad specificity towards different RES molecules tested. The results suggest a link between OPR3, GSH, and ASC and a function in RES/ROS-detoxification.

## Results

### OPR3 possesses a wide substrate spectrum

The OPR3 coding sequence from Arabidopsis was cloned from total leaf RNA, and the protein heterologously expressed in *E. coli* as previously described by Maynard et al. (2018a). As revealed by SDS-PAGE and Coomassie Brilliant Blue staining, isolation and purification yielded a major band constituting about 90% of overall protein with an apparent molecular mass of 42 kDa in good agreement with the calculated Mr of 42.69 kDa (Q9FUP0) (Fig. 1A). The old yellow enzyme OPR3 contains FMN as cofactor. Photometric analysis of purified OPR3 confirmed the presence of FMN. The absorption spectra (Fig. 1A) of OPR3 revealed absorption maxima at 280 nm, 374 nm and 467 nm, compared to free FMN at 267 nm, 374 nm and 445 nm, which was similar to results obtained from other OYE’s (Brown et al. 1998). Previous studies indicated that OYEs catalyze the oxidation of a wide range of ‘ene’ compounds (Williams and Bruce 2002; Toogood et al. 2010; Toogood and Scrutton 2014). As shown for OPR3 from tomato and some other species, this broad substrate specificity seems to be a common feature of OPR3 enzymes (Schaller et al. 1998; Straßner et al. 1999; Costa et al. 2000; Maynard et al. 2018b).

**Fig. 1.**
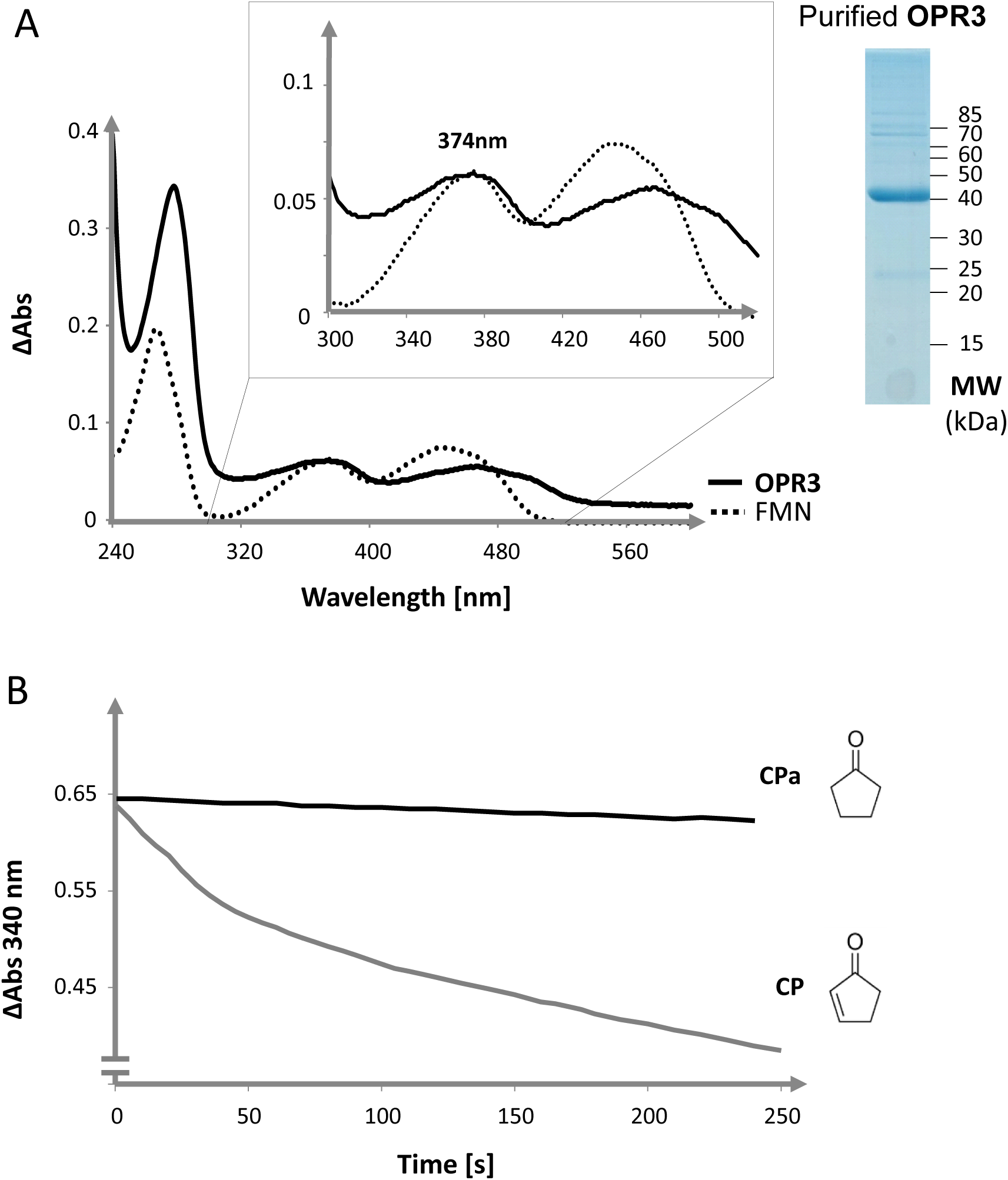
Purification of recombinant OPR3 and activity of OPR3 with cyclopentenone as substrate. (A) UV-Vis spectrum and Coomassie Brilliant Blue-stained electropherogram of heterologously expressed OPR3 (20 µg) revealed successful purification of the holoprotein (purity 90%) with the apparent molecular mass of 42 kDa and absorption maxima appearing at 278, 374 and 467 nm. Presence of FMN in OPR3 was confirmed by spectral comparison of free FMN (dashed line) and OPR3 (bold line). The inset highlights spectral overlap at 374 nm. (B) NADPH oxidation activity of 0.1 µmol/L recombinant OPR3 protein in the presence of 570 µmol/L cyclopentenone (CP) or cyclopentanone (CPa) and 150 µmol/L NADPH. The specificity of OPR3 spectra and activity was confirmed through empty vector control(s).

To provide an overview on the substrate specificity of OPR3 and to address possible *in vivo* functions, several potential substrates were tested with emphasis on detoxification of RES molecules which are physiologically relevant in *planta*. Starting point was 2-cyclopentene-1-one which may be considered as minimal reactive electrophilic unit of non-enzymatically and enzymatically formed cyclopentenones (Fig. 1B). The addition of 2-cyclopentene-1-one (CP) to an assay containing OPR3 and NADPH, led to a decrease in absorption at 340 nm. This was not the case when assayed with its homologue cyclopentanone (CPa), revealing that saturation of the α,β-unsaturated bond of CP occurred. Further tests revealed that the RES model compound N-ethyl-maleimide (NEM) is an efficient substrate of OPR3. Table 1 assembles the results for all tested substrates where OPR3 activity was evident. Methyl vinyl ketone (MVK) and 1, 4-benzoquinone (BQ), which is the reactive moiety of reaction products of ROS with tocopherol (Munné-Bosch et al. 2007; Deller et al. 2008), and quercetin (Boots et al. 2005) were also easily accepted substrates of OPR3 in line with results obtained for other OYEs (Vaz et al. 1995 and references cited herein). Further tests revealed that in contrast to the α,β-unsaturated cyclic and aliphatic ketones, the reactive aldehyde malondialdehyde, quantified in peroxisomes by Palma et al. (1991), was not a substrate for OPR3.

**Table 1.**
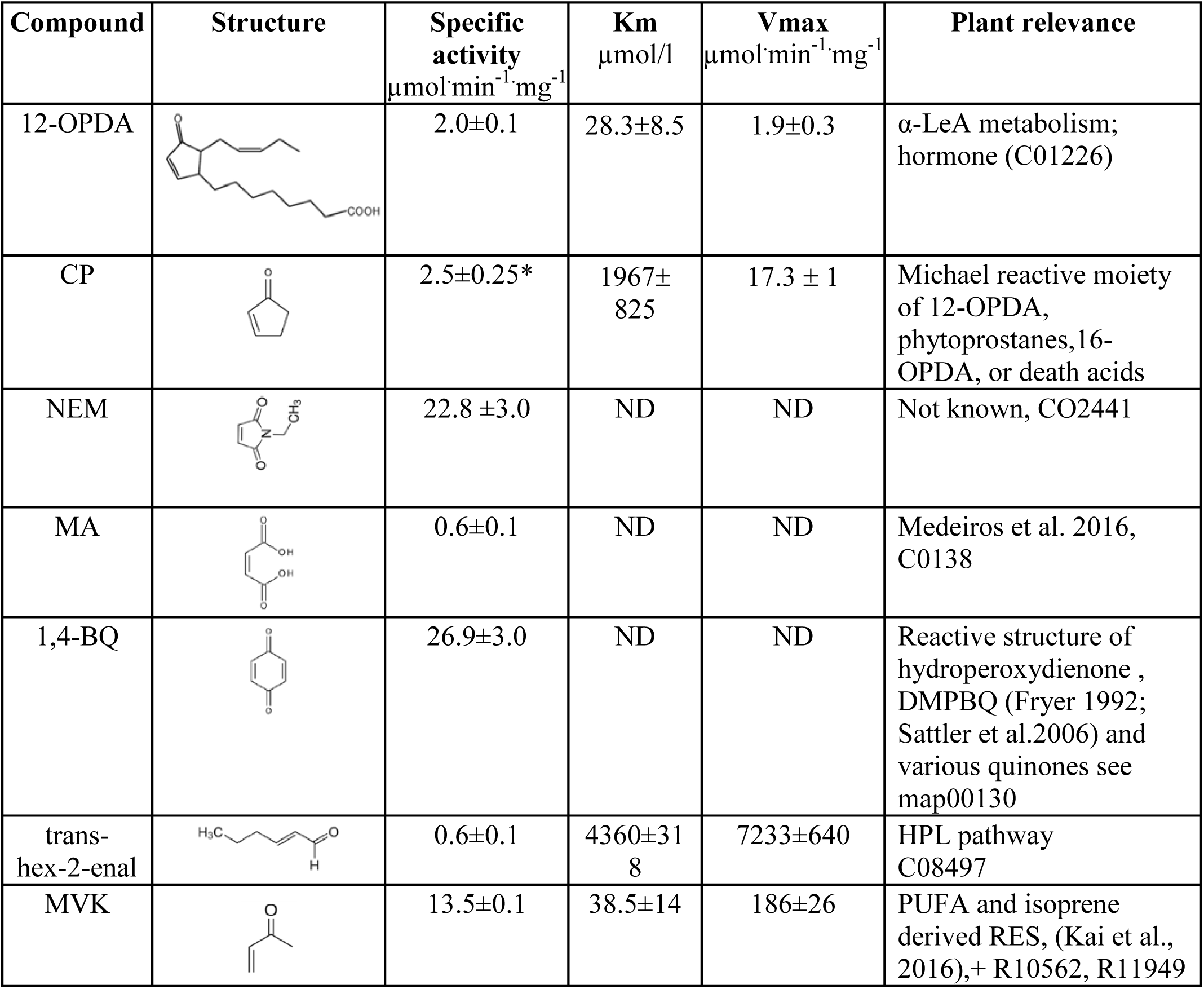
Kinetic parameters of OPR3 for the reduction of tested electrophiles.

| Compound | Structure | Specific activity<br>$\mu\text{mol}\cdot\text{min}^{-1}\cdot\text{mg}^{-1}$ | Km<br>$\mu\text{mol/l}$ | Vmax<br>$\mu\text{mol}\cdot\text{min}^{-1}\cdot\text{mg}^{-1}$ | Plant relevance |
| --- | --- | --- | --- | --- | --- |
| 12-OPDA | | 2.0±0.1 | 28.3±8.5 | 1.9±0.3 | α-LeA metabolism; hormone (C01226) |
| CP |  | 2.5±0.25* | 1967±825 | 17.3 ± 1 | Michael reactive moiety of 12-OPDA, phytoprostanes, 16-OPDA, or death acids |
| NEM |  | 22.8 ±3.0 | ND | ND | Not known, CO2441 |
| MA |  | 0.6±0.1 | ND | ND | Medeiros et al. 2016, C0138 |
| 1,4-BQ |  | 26.9±3.0 | ND | ND | Reactive structure of hydroperoxydienone , DMPBQ (Fryer 1992; Sattler et al.2006) and various quinones see map00130 |
| trans-hex-2-enal |  | 0.6±0.1 | 4360±318 | 7233±640 | HPL pathway C08497 |
| MVK |  | 13.5±0.1 | 38.5±14 | 186±26 | PUFA and isoprene derived RES, (Kai et al., 2016),+ R10562, R11949 |

Two ‘ene’-containing dicarboxylic acids were tested, namely maleic acid (MA) and traumatic acid (TA). Similar to the results obtained for the OPR3 orthologue from tomato (Straßner et al. 1999), MA was a poor electron acceptor with low specific activity of 0.6 µmol.min^-1^.mg^-1^. Further, in contrast to the LOX-hydroperoxide lyase (HPL) pathway metabolite TA, the HPL-derived oxylipin trans-hex-2-enal served as substrate for OPR3 with a K_m_ of 4.36 mmol/L. OPR3-mediated conversion was undetectable with the peroxisomal aromatic ‘ene’-compounds trans-cinnamic acid and trans-cinnamic-aldehyde (Kao et al. 2018). Cis-isomers could not be tested due to the lack of commercial availability.

### OPR3 efficiently protects thiols by reducing RES

Thiol reactivity is a common feature of RES. OPR3 detoxifies thiol-reactive RES and, thereby, may protect thiols. The efficiency of OPR3-mediated RES sequestration was indirectly assessed by incubating GSH with NEM at various stoichiometric ratios with or without OPR3. Subsequently, remaining free thiols of GSH were titrated with DTNB. NEM blocks the free thiols of GSH, which then cannot react with DTNB anymore. As expected, samples containing GSH (200 µmol/L) and increasing amounts of NEM in the presence of NADPH (300 µmol/L) but without OPR3 lost higher proportions of free thiols (Fig. 2, lower curve). In a converse manner the presence of OPR3 (0.27 µmol/L) almost completely protected GSH from alkylation (Fig. 2, upper trace). The *in vitro*-results demonstrate the ability of OPR3 to inactivate NEM as Michael acceptor by reducing its double bond (chemical structure see Table 1) and is in line with the hypothesized role of OPR3 in protecting low molecular weight antioxidants *in vivo*.

**Fig. 2.**
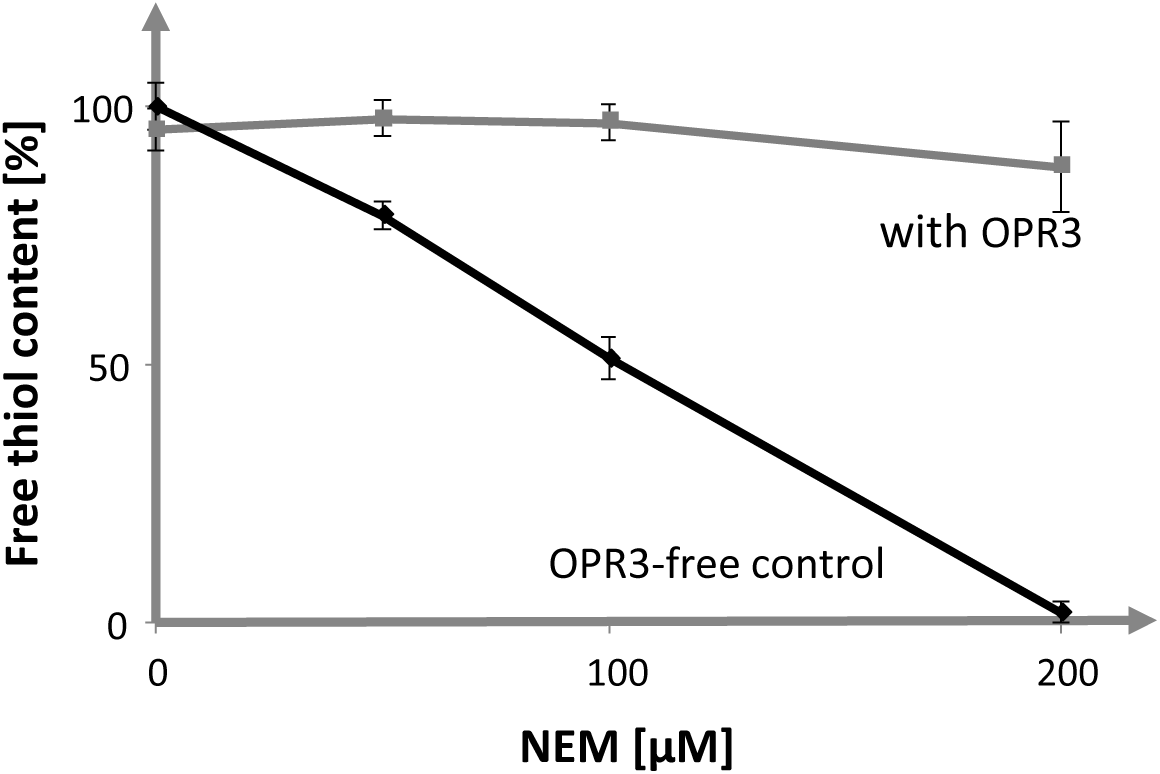
Protection of GSH from reacting with NEM in the presence of OPR3. GSH (200 µmol/L) was pre-incubated in the presence of different concentrations of NEM (0-200 µmol/L), with or without OPR3. The ratio of 1 relates to 200 µmol/L GSH incubated with 200 µmol/L NEM ± OPR3. Values represent means±SD (n=12) and are expressed as percentage of the maximum thiol content (100 %=196.6±1.7 µmol/L free thiols). For details see Materials and Methods.

Enzymatic RES scavenging may be particularly important if ascorbate and glutathione are low, e.g. as in seeds (Pena-Ahumada et al. 2006) and in leaves of stressed plants. *In silico* expression analysis (https://www.arabidopsis.org/) revealed that *OPR3* transcript accumulation under abiotic stress treatments, such as osmotic, salt and drought stress, correlates with that for typical RES scavenging enzymes like chloroplastic aldo-keto reductase (At2g37770) or NADPH-dependent aldehyde reductases (At1g54870 and At3g04000) in shoots or roots of Arabidopsis (Yamauchi et al. 2010).

To address the physiological role of OPR3 in detoxifying RES, we performed Arabidopsis leaf disc assays. Leaf discs of WT (col-0) and *opr3* (col-0) plants were exposed to MVK or 12-OPDA at a concentration of 0.5 mmol/L. Relative to WT plants we expected that *opr3* mutants are more susceptible to RES-induced plant damage in terms of PSII efficiency. Besides severe time-dependent effects of MVK and 12-OPDA on the PSII yield, no significant difference between the *opr3* mutant and WT plant was evident (Fig. 3).

**Fig. 3.**
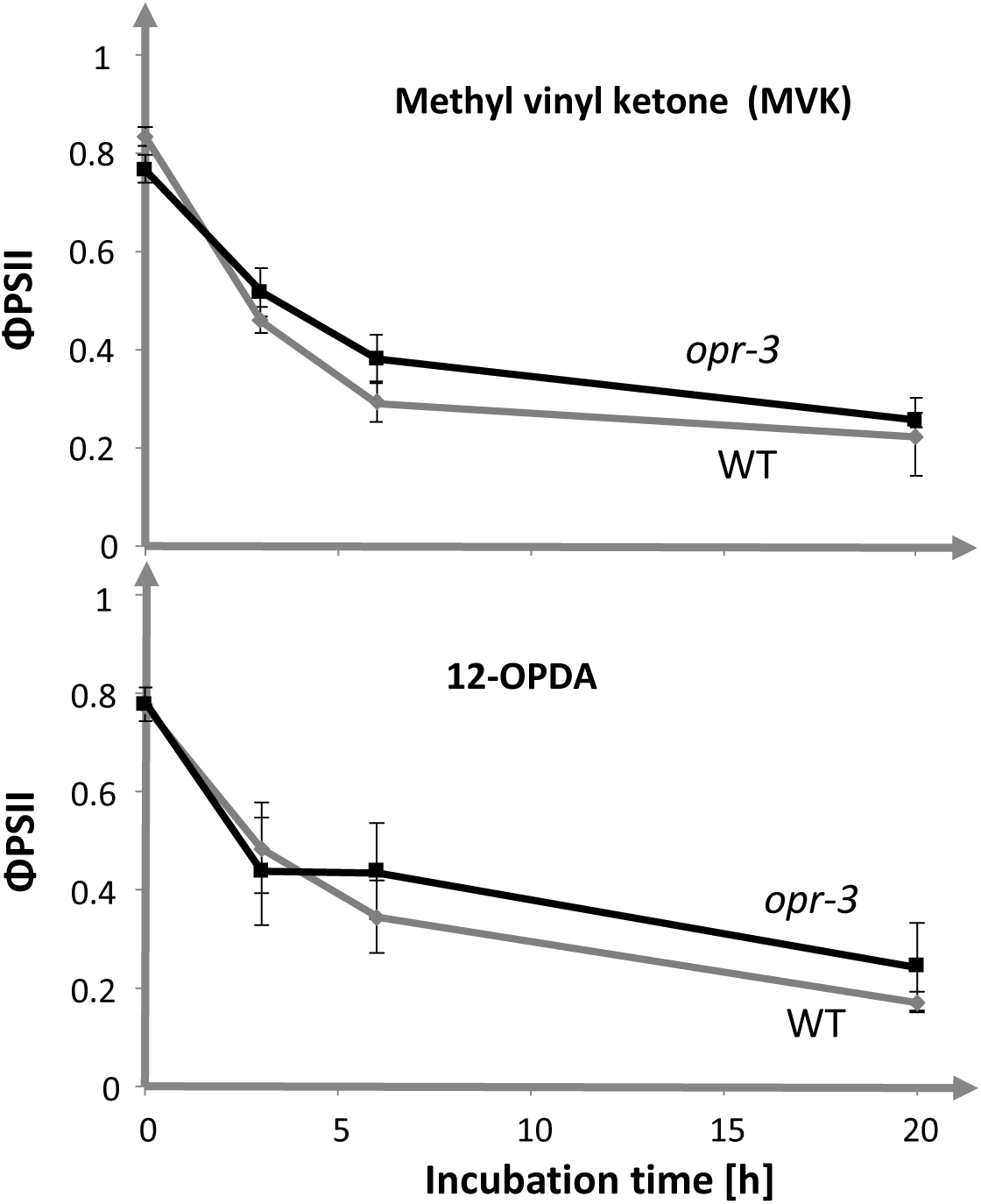
Changes in quantum yield of photosystem II (ΦPSII) of WT and *opr3* in response to treatment with 12-OPDA and methyl vinyl ketone (MVK). Leaf discs of WT (Col-0) and *opr3* (Col-0 background) floated on solution containing the indicated compounds (0 or 5 mmol/L) in 100 µmol/L CaCl_2_. ΦPSII was determined by chlorophyll a fluorescence analysis using the pulse-amplitude-modulated photosynthesis yield analyzer (Mini-PAM). Data are means±SD (n=6).

### Potential role of ASC in OPR3 activity regulation

In order to further characterize the broad substrate specificity of OPR3, we queried literature for properties of related flavoenzymes like cyclopentanone-1,2-monooxygenase (1.14.13.16). Cyclopentanone-1,2-monooxygenase enzymatically forms lactones from cyclopentanones. Although OPR3 does not form lactones itself, the affinity to the cyclopentenone moiety witnessed through its substrate 12-OPDA directed our attention to the possibility that abundant peroxisomal lactones like ASC might affect OPR3 activity. This hypothesis was addressed by investigating the possible interaction of ASC with OPR3. ASC quenched the tryptophan (Trp) fluorescence of OPR3 like the positive controls 12-OPDA and NEM, albeit with lower efficiency (Fig. 4A). Half-maximal Trp fluorescence quenching by ASC occurred at a concentration of 1.7 mmol/L (Fig. 4B). The results suggest that ASC or short lived species such as monodehydroascorbate (MDHA) (Du et al. 2012) may interact with OPR3 in the peroxisome.

**Fig. 4.**
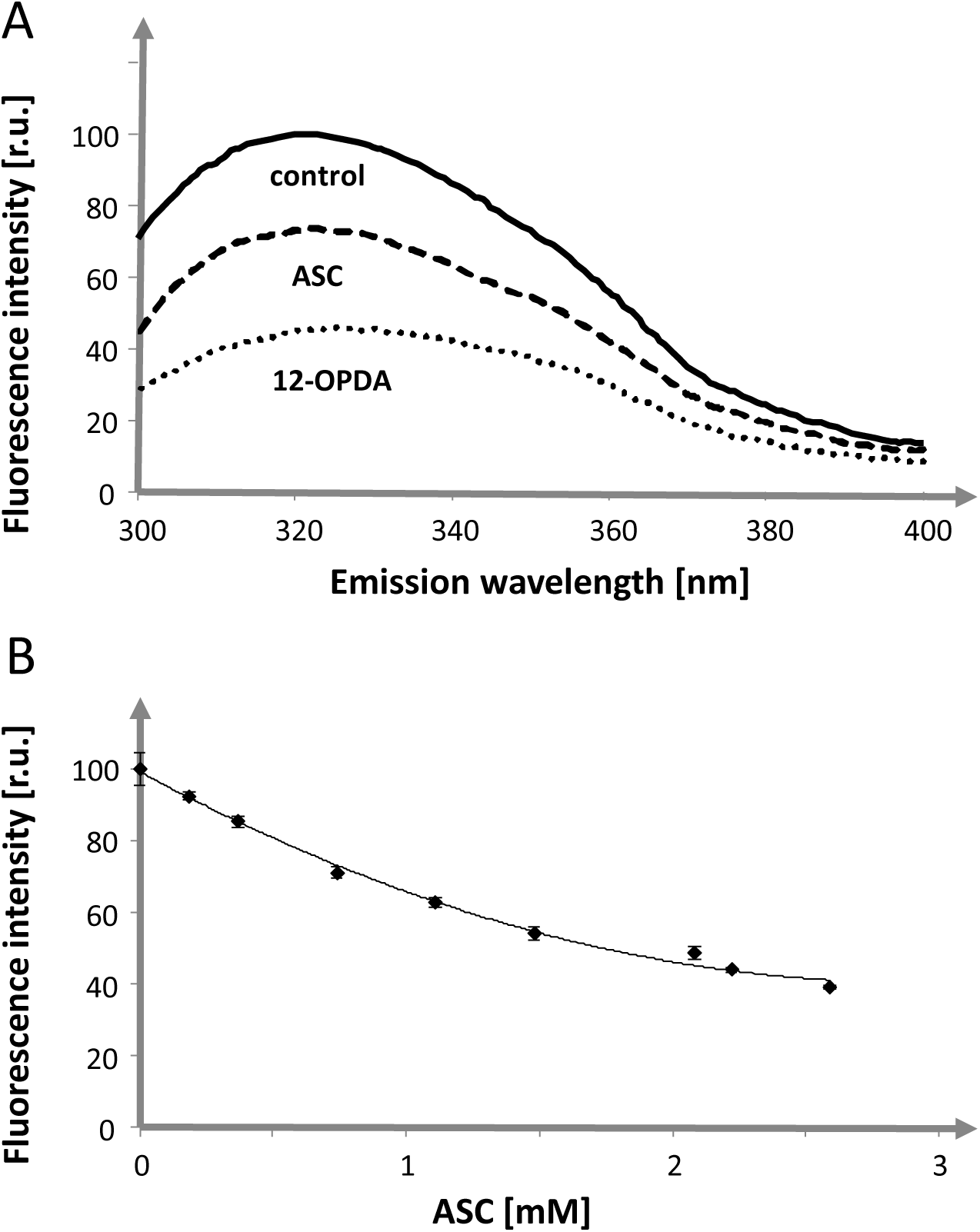
Effect of ascorbate (ASC) on tryptophan fluorescence of OPR3. (A) Interaction of OPR3 with ASC as indicated by changes in intrinsic Trp-fluorescence in comparison to 12-OPDA as positive control. ASC and 12-OPDA at a final concentration of 607 µmol/L quenched the intrinsic Trp-fluorescence of 0.75 µmol/L OPR3 by 26.0±1.3% and 54.0±0.4% in comparison to control buffer (100.0±8.3%). Data are means±SD, n=3. (B) Effect of increasing ASC concentration on intrinsic Trp fluorescence quenching of OPR3. The decline in relative fluorescence intensity (RFI) depended on ASC concentration. Fitting a polynomial equation of 2^nd^ order, y=6.901x^2^-40.309x+99,247, R^2^= 0.9947 revealed that 50% Trp quenching of OPR3 occurred at 1.7 mmol/L ASC. Data are means±SD, n=3.

Peroxisomal enzymes like catalase and ascorbate peroxidase are inhibited by their ligands (Durner and Klessig 1995). However the enzyme activity remained unaltered when ASC was added to the OPR3 activity assay in the range of 0.5-2 µg total enzyme and 1-5 mmol/L ASC (data not shown). Another possibility that seemed likely was the interaction of ASC with peroxisomal substrates of OPR3. Fodor et al. (1983) reported adduct formation between ASC and MVK under physiological conditions. Therefore, we incubated 12-OPDA with ASC in phosphate buffer, pH 7.2, at 25°C for 3 or 6 h and assayed for OPR3 activity. As a control we incubated 12-OPDA with GSH, expecting conversion of the OPR3-substrate OPDA to the OPDA-GSH adduct and therefore decreased OPR3 activity (Fig. 5A).

**Fig. 5.**
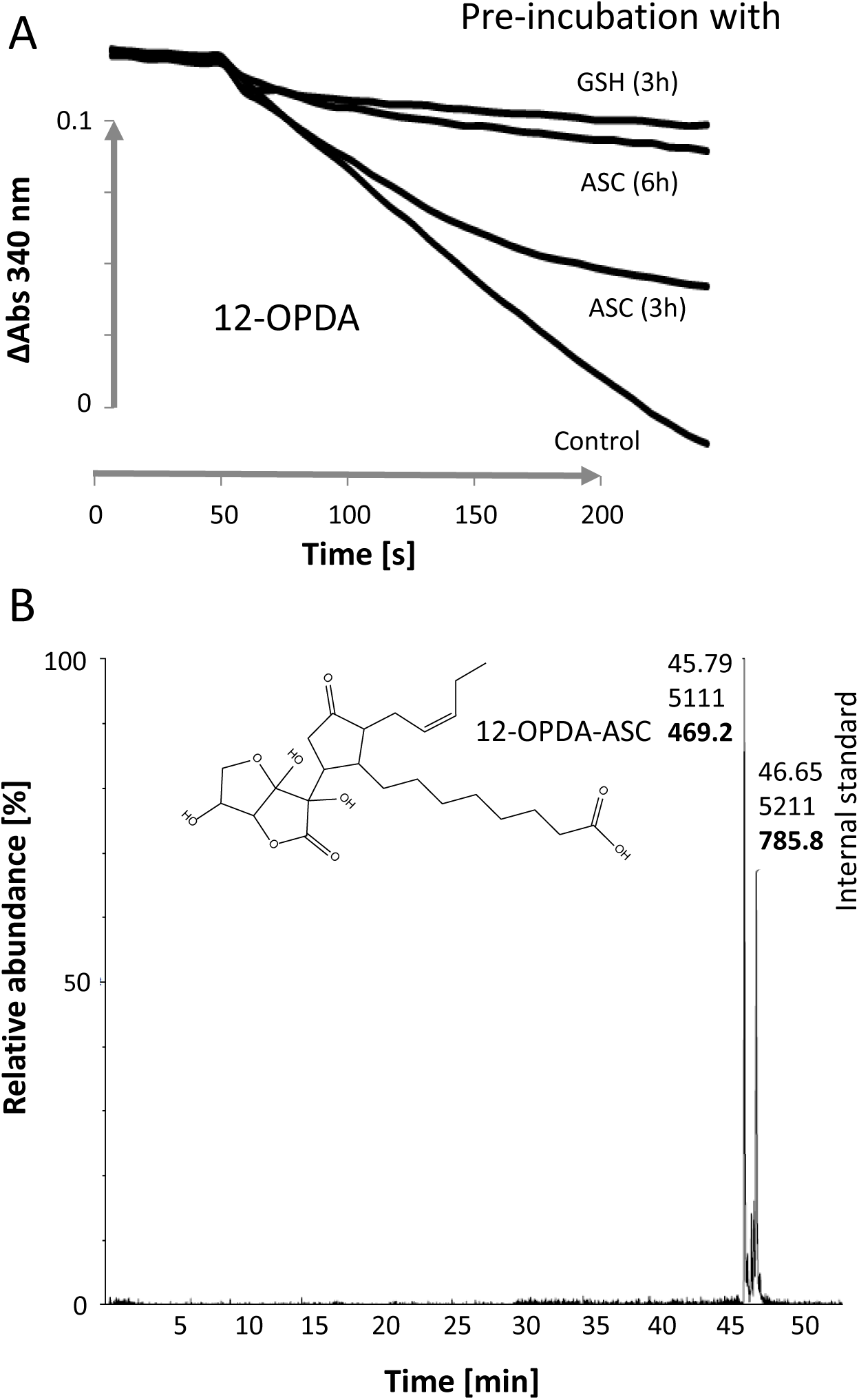
Modification of OPR3 substrates by ascorbate and glutathione. (A) OPR3 activity with 12-OPDA pre-incubated with GSH or ASC. 12-OPDA was pre-incubated with antioxidants at molar ratio of 2.5 for GSH and 20 for ASC for indicated time intervals. The experiment was carried out three times, and representative curves are shown. (B) Nano-UPLC chromatogram of 12-OPDA incubated with ASC. The MS analysis of the peak eluting at 45.8 min revealed that 12-OPDA covalently interacts with ASC yielding 12-OPDA-ASC (m/z=469.2431 [M+H]^+^-Ion) (see supplemental Figures 3 and 4). The predicted structural formula is shown. Also highlighted is the retention time of the lock mass compound GluFib, [M+2H]^2+^, m/z 785.8418 (Suppl. Figures 3 and 4).

After pre-incubation of 2 mmol/L 12-OPDA with 40 mmol/L ASC for 3h, OPR 3 activity was reduced by 28±5% (n=3) compared to 12-OPDA alone. The positive control included GSH (5 mmol/L) which is known to modify 12-OPDA (2 mmol/L) and inhibited OPR3 activity in our test system. After preincubation of 12-OPDA with ASC for 6 h, the inhibition of turnover was similar as in the case of GSH/12-OPDA-preincubation indicating that the “ene” unit of 12-OPDA was covalently modified by ASC (Fig. 5A). Isothermal titration microcalorimetry (ITC) revealed that the interaction proceeds at equimolar ratio and as exothermic reaction (Suppl. Fig. 1). Analysis of the reaction products by nano-UPLC/nano-ESI-MS proved the formation of the covalent adduct of 12-OPDA with ASC. The novel compound was detected at a retention time of 45.8 min with an *m/z* value of 469.2431 (*m/z*_theo_ 469.2432, Δ=0.23 ppm, Fig. 5B, Suppl. Fig. 2-4). An additional signal appeared in the mass spectra, which corresponded to a single water loss (m/z 451.2322) of 12-OPDA-ASC. This water loss may either be the result of a fragmentation of 12-OPDA-ASC in the ion source, or a condensation product of 12-OPDA and ASC was formed during the incubation.

### OPR3 has monodehydroascorbate reductase activity and regenerates ASC

The relationship between ASC and OPR3 was scrutinized further by *in silico* analysis. We focused on ASC interacting enzymes and revealed that the monodehydroascorbate reductase (MDHAR) belongs to the same class of oxidoreductase enzyme as the oxophytodienoic acid reductases. Despite their low sequence identity of less than 10%, both reductases bind NADPH and flavo-cofactors and structurally arrange into alpha-beta domains. The structural superposition of OPR3 with cytosolic MDHAR from rice Os09g0567300 (Fig. 6A) revealed coinciding positioning of the cofactor binding sites in both oxidoreductases. In addition ASC was positioned in the wide substrate binding pocket of OPR3 in close proximity (<0.5 nm) to the cofactor (Fig. 6B). The model supports the proposed ability of OPR3 to interact with ASC in line with our results from Trp fluorescence analysis (Fig. 4).

**Fig. 6.**
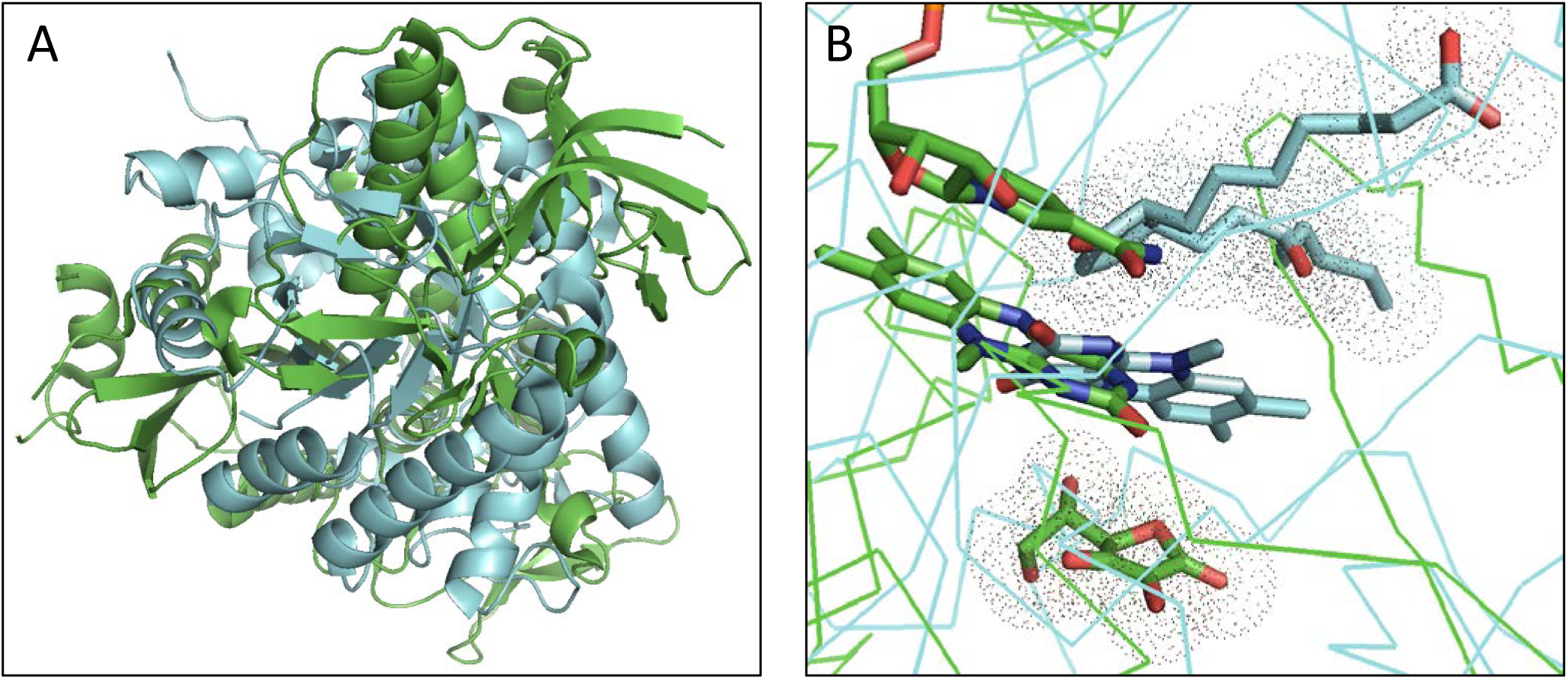
Comparison of the protein structures of OPR3 and monodehydroascorbate reductase (MDHAR). Structural superposition of both enzymes using PyMOL (RMS of 22.55). (A) Cyan and green 3-D cartoon represent the OPR3 (2g5w) and MDHAR (5jcn) crystal structures, respectively. (B) Structural alignment highlighting the ASC molecule bound to MDHAR (green) and the 12-OPDA homologue 8-iso PGA_1_ bound to OPR3 (cyan). In addition, the flavin cofactors are shown revealing that the same binding cleft is occupied by the cofactors in both enzymes. The NAD bound to MDHAR is partially depicted, for clarity, structure is presented in ribbons, and the isoalloxazine rings are shown for the cofactors FAD and FMN. The atoms in highlighted molecules are depicted as sticks and are colored as follows: green, carbons of ascorbate, NAD and FAD; cyan, carbons of 8-iso PGA1 and FMN; blue, nitrogen; red, oxygen.

The possibility that OPR3 functions as MDHA reductase was addressed in an MDHAR activity test (Hossain et al. 1984, Park et al. 2016). Ascorbate oxidase (AO) was added to ASC to generate MDHA. The absorbance at 340 nm decreased rapidly with time in the presence of OPR3, ASC, NADPH and AO (Fig. 7A), demonstrating that OPR3 regenerated ASC from *in situ* formed MDHA. NADPH-dependent reduction of *in situ* formed MDHA by OPR3 was characterized by measuring NADPH oxidation at various steady state concentrations of MDHA, generated by applying different AO units to fixed ASC amount (7B). This approach yielded an apparent K_m_ of 1 µmol/L, in line with MDHAR from spinach (Hossain et al. 1984). The typical MDHAR substrate K_3_[Fe(CN)_6_] (Hossain and Asada 1985) was also a good OPR3 substrate, similar to NEM (Supplementary Figure 6).

**Fig. 7.**
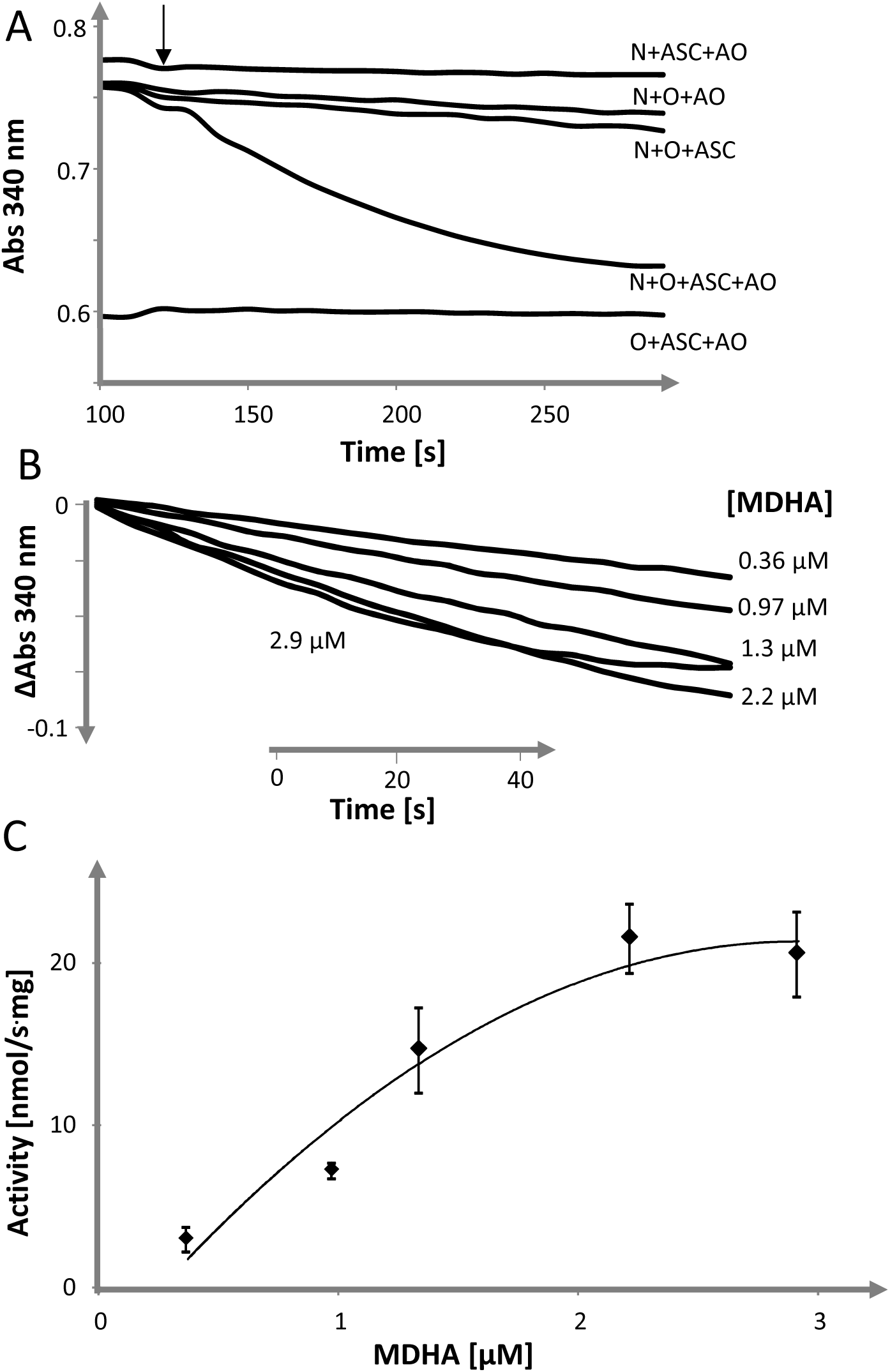
The MDHAR activity of OPR3. (A) 140 µl buffer (KPi, 40 mmol/L; pH 7.2) containing 1.1 µg OPR3 (O), 129 µmol/L NADPH (N) and 10.7 mmol/L ASC was incubated and 1 µl of 0.05 U ascorbate oxidase (AO, indicated with an arrow) was added after 120 s (N+O+ASC+AO). Several control reactions were recorded as well. (B) Representative graphs for determination of Michaelis Menten parameters of MDHAR activity of OPR3. OPR3 (0.5 µg) was incubated with 125 µmol/L NADPH, 1 mmol/L ASC and varying amounts of AO. The indicated MDHA concentrations were calculated in a separate experiment, (C) Michaelis Menten plot of OPR3 with MDHA as substrate was analysed by plotting steady state MDHA concentrations against initial velocities (20 s) of NADPH consumption exemplified in (B). Data are means±SD, n=3. For details see Materials and Methods.

We then analyzed *opr3* mutant and WT plants in a gas environment containing either ambient CO_2_ or very low CO_2_, enhancing photorespiratory H_2_O_2_ release by glycolate oxidation in the peroxisomes. This condition aimed for challenging the involvement of OPR3 in ASC regeneration *in vivo* after 3 and 6 weeks of germination (Fig. 8). ASC and dehydroascorbate (DHA) amounts were quantified in leaf extracts and revealed that *opr3* compared to WT displayed lower quantities of reduced ascorbate. The ratio of DHA/ASC was significantly higher in 6 week old *opr3* than in WT plants, both in 460 ppm and 37 ppm CO_2_ (two tailed Student’s T-test with uneven variance, p<0.1), but not between the treatments, and when combining the results from both regimes with p<0.01. The decreased amounts of reduced ASC and the increased DHA/ASC-ratios in *opr3* plants underline our *in vitro* observations and tentatively support the hypothesis that OPR3 functions in the regeneration of the radical scavenger ASC in peroxisomes (Fig. 8).

**Fig. 8.**
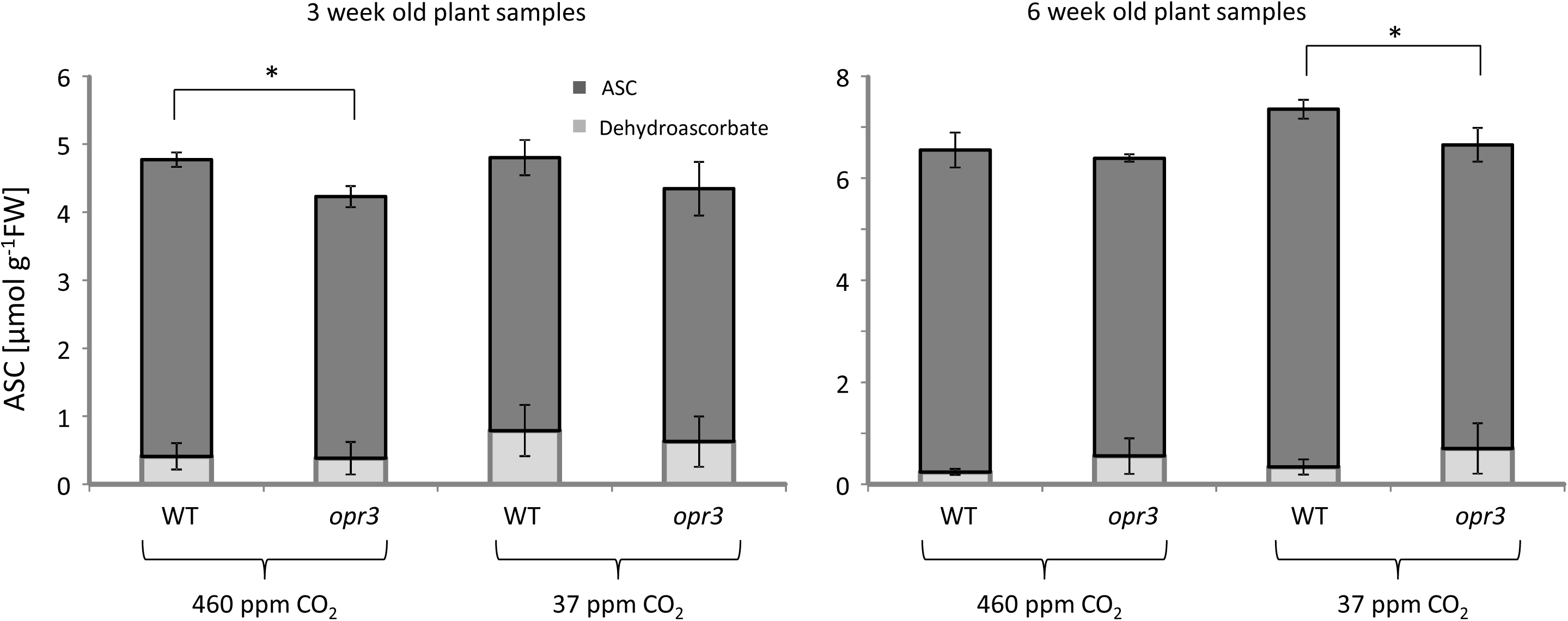
Ascorbate and dehydroascorbate contents of WT and *opr3* plants under photorespiratory conditions (37 ppm CO_2_). Three and six weeks old WT and *opr3* plants were flushed with low CO_2_-air. Control plants were flushed with ambient air (460 ppm CO_2_, see Material and Methods). Significance is indicated as asterisks (p<0.05) and was calculated using Student’s two sided *t*-test assuming unequal variance.

## Discussion

Despite previous characterization of OPR3, a detailed description of substrates and ligands was lacking. Therefore, this study aimed to fill the gap in understanding OPR3 activity. The results revealed that physiologically relevant RES and their derivatives are substrates of OPR3. It is interesting that among the tested RES molecules, trans-hex-2-enal, a major product of LOX multienzyme complexes was recognized as substrate of OPR3. However, the low affinity of OPR3 toward trans-hex-2-enal contrasts the high affinities of aldo-keto reductases previously implicated in the modulation of green leaf volatiles with K_m_-values in the µmol/L range (Mano et al. 2002; Tanaka et al. 2018). The data suggest that under physiological conditions trans-hex-2-enal is probably not a substrate of OPR3 or only a minor substrate. Instead, the cytosolic NADPH-dependent oxidoreductase 2-alkenal reductase (AER; EC 1.3.1.74) was reported by Mano et al. (2005) to efficiently reduce trans-hex-2-enal. These authors used chlorophyll fluorescence as the readout of RES-induced stress in plants and showed that overexpression of AER in the cytosol protects cells form reactive carbonyls. The contribution of OPR3-dependent RES detoxification in cellular RES management will require deeper analysis of peroxisome-related parameter(s).

OPR3 oxidizes NADPH in the presence of CP, the RES moiety of phytoprostanes and 12-OPDA (Mueller et al. 2008) (Fig. 1). This observation is consistent with the affinity of OPR3 for cyclic “enones” such as cyclohexenone (Costa et al. 2000), phytoprostanes (Mueller et al. 2008) or OCPD (Maynard et al. 2018a). The first description of the α-LeA-derived metabolite 12-OPDA as substrate of OPR3 occurred by Vick and Zimmerman in 1986. The reported K_m_ values ranged between 14 and 190 µmol/L. The 14-fold variation might be due to varying purity of 12-OPDA as only the cis-form is efficiently reduced by OPR3, and the inconsistent NADPH concentrations utilized in the assays.

The determined K_m_ value of OPR3 for 12-OPDA was 28.3 µmol/L and thus in line with published data. Similar to tomato OPR3 (Straßner et al. 1999), NEM was also efficiently reduced by OPR3 (specific activity of 22.8 ±3.0 µmol.min^-1^.mg^-1^) and as demonstrated in Fig. 2, OPR3 protected thiols from reacting with NEM. The dione analogue of NEM, 1,4-BQ might be a physiological relevant substrate for OPR3, as it is the reactive moiety of highly unstable intermediates of oxidized tocopherol and quercetin (Table 1). Fryer et al. (1992) showed that oxidation products of tocopherols are formed by reaction of tocopherols with oxidized α-LeA. Thus there exists a link between 1,4-BQ and the oxylipin pathway. At present it is not clear whether the 1,4-BQ reduction proceeds via one electron transfer as described for some quinone reductase enzymes (see Deller et al. 2007) or directly by two electron transfer as known for NADPH-dependent OYE (Vaz et al. 1995). Exposure of leaf tissue to RES leads to malfunctioning photosynthesis and, if available, the RES-detoxifying enzymes prevent plants from PSII damage (Alméras et al. 2003, Tanaka et al. 2018). The photosynthetic performances of *A. thaliana* WT (Col-0) and *opr3* (Col-0) plants exposed to the highly affine OPR3 substrates MVK and 12-OPDA (Fig. 3) revealed no differences in their sensitivity towards these RES-molecules, possibly indicating either unaltered detoxification potential of the plants or involvement of other sub-cellular factors. Although this result questions the physiological role of OPR3 in RES detoxification, subcellular metabolite analysis would be needed to scrutinize this dependency further and resolve the associated mechanisms. Small α,β-unsaturated carbonyl compounds as acrolein and methyl vinyl ketone administered to leaves have been reported to enhance expression of the pathogenesis-related gene PR4. Thus there certainly is a subcellular perspective in detoxification, because OPR3 is localized in the peroxisome (Alméras et al. 2003). In this context it would be interesting to explore the substrate spectrum of the cytosolic OPR2.

The unaltered sensitivity to tested RES-molecules including OPDA in *opr3* mutants compared to WT may be caused by enhanced GSH synthesis stimulated by accumulating OPDA as shown in feeding experiments (Park et al. 2013). Also, RES-induced upregulation of transcripts encoding other OPR-isoforms (reductases) like *OPR1*, as reported in Arabidopsis in response to MVK, could also contribute in RES-detoxification (Alméras et al. 2003). Similarly, transcripts of *OPR2* were shown to be upregulated in response to OPDA-exposure, while glutathione metabolism-related enzymes were part of the ORGs (Taki et al. 2005). Further analysis will be required to explore, identify, and characterize the function of OPR3 as alternative NADPH-dependent reductase in *planta*.

Comparisons of cofactor-binding domains can unveil functional similarities at the structural level even for proteins with low sequence similarities (Mascotti et al. 2016). The inspection of the structure of flavin-dependent oxidoreductases and their substrates indicated that OPR3, cyclopentanone-1,2-monooxygenase, and MDHAR are distantly related. The similarity of the cofactor binding domains of OPR3 with MDHAR suggests that OPR3 binds ASC in the wide substrate binding pocket in vicinity of the cofactor. Our *in vitro* studies also revealed that ascorbate interacts with OPR3. More importantly, the oxidation product of ASC generated by ascorbate oxidase served as substrate for OPR3, a feature shared with MDHAR located in peroxisomes and cytosol. Similar to spinach MDHAR (Hossain et al., 1984), the Km value of OPR3 for MDHA was determined with 1µmol/L. Although the MDHAR enzymes from peroxisomes have been characterized from Arabidopsis, no Km values were determined (Lisenbee et al. 2005). The relative contribution of peroxisomal MDHAR, 2 isoforms annotated at present, with one yet unidentified MDHAR see Lisensbee et al. (2005) and OPR3 in MDHA reduction *in vivo* awaits clarification.

The MDHA reduction mechanism of OPR3 does not involve NADPH-coupled oxygen consumption (see supplementary Figure S6). Also, superoxide dismutase (SOD) supplementation of the reaction mix did not inhibit MDHAR activity of OPR3 indicating that superoxide radicals, formed through OPR3-mediated electron transfer from NADPH to oxygen, are not responsible for MDHAR activity of OPR3 (data not shown) opposite to the superoxide anion-mediated MDHAR activity of mitochondrial apoptosis-inducing factor (Miramar et al., 2001).

The MDHAR activity of OPR3 may be important under metabolic conditions where ASC is rapidly oxidized in ascorbate peroxidase-dependent H_2_O_2_ detoxification. The physiological relevance of the two here proposed MDHA-reducing activities in ASC-regeneration of peroxisomes may be linked to distinct regulation or sensitivity to inhibition and turnover. Co-expression analysis (calculated with ATTED-II, [http://atted.jp/]) for OPR3 showed that homologues of OPR3 co-express with MDHAR3 (At3g09940) and MDHAR2 (At5g03630) in rice. Likewise, analysis of Arabidopsis genes co-expressed with OPR3 and MDHAR3 revealed a high overlap (data see supplementary table, Table supplement S1). These results suggest either irreplaceable redundancy or unique complementarity of OPR3 and MDHAR function, in the first case to realize the maximal detoxification capacity, in the latter case by distinct features that still have to be explored. OPR3 and peroxisomal MDHAR (At3g52880) are highly abundant leaf proteins ranking in the group of the top 5-10% most abundant polypeptides (345^th^ position for OPR3) and 181^th^ for MDHAR [At3g52880] out of 4190 polypeptides), while peroxisomal MDHAR4 (At3g27820) is less abundant (2151^th^ out of 4190 polypeptides). Thus it will be interesting to quantify the contribution of OPR3-dependent MDHA reduction capacity relative to MDHAR activity in the peroxisome in future work.

On the basis of results described in this study, we suggest that apart from its function in converting 12-OPDA into JA precursors, OPR3 plays additional roles in RES-detoxification as well as in reduction of MDHA to ASC. Bi-functionality is not unusual for enzymes localized in peroxisomes as exemplified by firefly luciferase or the transthyretin-like (TTL) protein (Oba et al. 2003; Lamberto et al. 2010) which likely is due to the multifaceted roles of peroxisomes in ROS regulation, metabolic conversions and inter-organelle communication (Kao et al. 2018). Our finding that 12-OPDA is covalently modified by ASC adds a novel adduct of this RES molecule to plant physiology. The corresponding GSH-12-OPDA adduct was characterized by Davoine et al. (2005) and subsequently identified under physiological conditions (Davoine et al. 2006). The physiological occurrence and bioactivity of ASC-12-OPDA should be investigated in future studies. If 12-OPDA-ASC is found, it will be interesting to determine its turnover and fate. The corresponding GSH-OPDA adduct is imported and cleaved in the vacuole (Ohkama-Ohtsu et al. 2010) similar to GSH-conjugates of xenobiotics (Wolf et al. 1996).

According to our proposed model (Fig. 9), OPR3 functions as MDHAR under oxidative stress thereby possibly competitively inhibiting 12-OPDA reduction and delaying or suppressing the expression of JRGs. The concomitant accumulation of 12-OPDA in turn would then stimulate GSH synthesis and transcriptionally activate antioxidant defense genes. The possibility of OPR isoforms acting as MDHAR and the potential of 12-OPDA and other RES to react with GSH and ASC add another level of complexity to the antioxidant defense system, and are issues which should be addressed further.

**Fig. 9.**
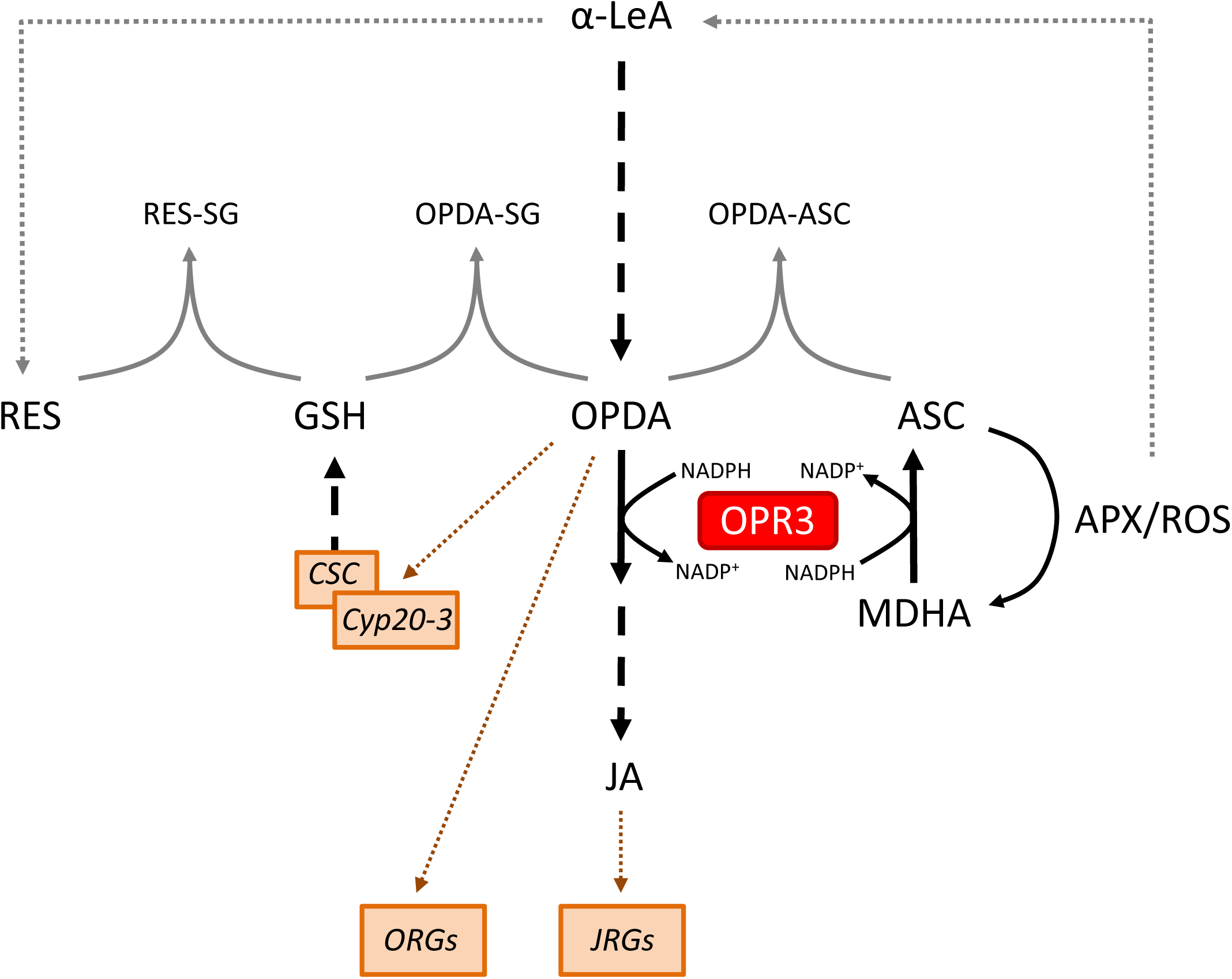
Proposed model of OPR3 being a bifunctional enzyme interconnecting ROS and RES-detoxification and signalling pathway. The simplified network is derived from results as demonstrated in this study and by literature survey mentioned in the text. The peroxisomal enzyme OPR3 interconnects the two major redox buffers GSH and ASC by NADPH dependent (1) degradation of 12-OPDA, which in turn influences GSH synthesis and (2) regeneration of reduced ASC from APX/ROS generated MDHA. In addition to 12-OPDA’s enzymatic transformation to JA precursor pools (bold broken line), 12-OPDA, as a RES compound is subject to chemical degradation by GSH in a feedback loop. In a similar fashion, OPR3 determines the fate of 12-OPDA through its conjugation with ASC, whose levels can be influenced by the MDHAR-like function of OPR3. (3) The reaction of ROS with α-LeA (dotted line) leads to diverse quenching products (i.e. RES) which may conjugate to these abundant redox buffers (shown for the thiol containing biomolecules like GSH) or are possibly prone to modification by ‘ene’ reductases, as has been addressed for OPR3 in this study. (4) Next to induced regulation (brown broken arrows) of GSH synthesis by 12-OPDA is the regulation of gene expression by 12-OPDA (ORGs) and JA (JRGs) that are part of this network, as they are involved in tocopherol, ASC α-LeA, ROS, RES and GSH metabolism.

## Materials and Methods

### Chemicals

Cyclopentenone, cyclopentanone, N-ethyl maleimide and 1,4-benzoquinone were purchased from Sigma-Aldrich (Darmstadt, Germany), traumatic acid from Tokyo Chemical Industry (Japan), MVK and trans-hex-2-enal from Acros-Chemicals (Thermo Fisher Scientific, Geel, Belgium). 12-OPDA was synthesized as described in Maynard et al. (2018a). 10 mmol/L and 20 mmol/L malondialdehyde stock solutions were generated from bis-dimethyl-acetal (Sigma Aldrich) in 1% H_2_SO_4_ as described in Esterbauer and Cheeseman (1990). All other chemicals were purchased from diverse commercial suppliers and were of highest purity. Ligands and substrates were prepared in 40 mmol/L potassium phosphate buffer (KPi) at pH values as indicated. The amount of ethanol (EtOH) or methanol (MeOH) was below 3% in the assays performed with chemicals dissolved in organic solvents.

### Synthesis of OPR3 and OPR3 activity tests

Cloning, expression and purification of OPR3 were performed as described in Maynard et al. (2018a). Enzyme purity was assessed by 12% SDS-PAGE and subsequent staining. Protein concentration was determined with Bio Rad assay. Photometric analysis of purified OPR3 (0.3 mg/ml in KPi 40 mmol/L pH 7.2) and phosphate (1µg/ml in KPi 40 mmol/L pH 7.2) was performed using a quartz cuvette and a Shimadzu spectrophotometer at medium scan speed, with a slit width of 2 nm and a sampling interval of 1 nm. The NADPH oxidation activity of the recombinant OPR3 protein with various substrates was measured aerobically at 25°C. OPR3 (0.5-3 µL, 0.24-0.66 mg/ml) was added to a cuvette containing 100 µL KPi 40 mmol/L, pH 7.2, and 15 µL NADPH (1.2 mmol/L) and placed in the Shimadzu spectrophotometer. After 1 min, 2 to 4.4 µL substrate was added and the decrease of absorption at 340 nm was recorded. The initial velocity within the first 0.5 min was taken for analysis. The velocity was linear for all assayed substrates except for CP, here only the slope of the first 40 s was linear. The affinity of OPR3 to NEM and 1,4-BQ was very high as the NADPH coupled absorption at 340 nm drastically decreased within 1 min. Michaelis Menten parameters were not determined (ND) for NEM, 1,4 BQ and MA. Specific activities were determined at 570 µmol/L concentration. Km and Vmax values were obtained from Lineweaver Burk and Michaelis Menten plots (trans-hex-2-enal). KEGG information (www.genome.jp/kegg/) and literature regarding physiological relevance of compounds or derivatives thereof in planta are provided in Table 1. NEM and MA were prepared in 40 mmol/L KPi pH 7.2 (buffer), 12-OPDA, 1,4-BQ, CP were dissolved in ethanol. Controls were conducted by omitting either enzyme, substrate or NADPH to ensure that the observed decrease in absorption was due to enzymatically coupled electron transfer from NADPH to substrate (Costa et al. 2000). The rates of NADPH oxidation (µmol NADPH.min^-1^.mg^-1^ protein) obtained for at least seven substrate concentrations were used to determine K_m_ and V_max_ values, using an extinction coefficient of 6220 M^-1^ cm^-1^. Data are means of at least three measurements ± SD.

The O_2_ consumption test of OPR3 in presence of substrate and NADPH was done as described for old yellow enzymes (Yamano et al. 1993). KPi-buffer (2.29 ml; 40 mmol/L, pH 7.2) containing 126 µmol/L NADPH and 742 µmol/L CP was incubated in a Clark type oxygen electrode (PC-operated Oxygraph+, Norfolk UK), at 50 rpm stirring speed. Detected O_2_ concentrations were in the range of 280±2.5 µmol/L and unchanged after injection of either 50 µL OPR3 (0.2 µg/µL) (n=3) or KPi 40 mmol/L pH 7.2 (n=3). O_2_ consumption was also not observed when OPR3 was incubated with NADPH before and after injection of CP (not shown).

MDHAR activity of OPR3 was assessed as described by Hossain et al. (1984). 120 µL buffer (KPi 40 mmol/L, pH 7.2) was mixed with 15 µL NADPH (1.2 mmol/L) and 2 µL OPR3 (0.57 mg/mL). After 120 s, 4.4 µL ASC (340 mmol/L in buffer) was added. To study the effect of AO-mediated MDHA generation and its successive reduction to ASC by OPR3, AO (Sigma Aldrich) was added and the decrease of NADPH was recorded at 340 nm. Controls ensuring that OPR3 catalyzed the recycling of ASC from AO-generated MDHA were performed as mentioned in text. The K_m_ value of OPR3 towards MDHA reduction was determined as follows. 0.5 µg OPR3 was incubated with 125 µmol/L NADPH, 1 mmol/L ASC, and varying amounts of AO (0.01875, 0.0375, 0.075, 0.15, and 0.21 Units) in a total volume of 145 µL with 50 mmol/L TRIS buffer, pH 7.8. By addition of AO, a decrease in absorbance at 340 nm occurred and the initial 20 s decrease in absorption due to NADPH consumption were considered for OPR3 catalysed reduction of *in situ* formed MDHA. The MDHA concentration in the enzymatic assay was determined photometrically at 360 nm in a separate experiment by variation of AO as above added to 1 mmol/L ASC, with 50 mmol/L TRIS; pH 7.8 buffer with the molar extinction coefficient (ε=3300 L.mol^-1^.cm^-1^) as described in Hossain et al. (1984). Measurements were performed in triplicate.

### OPR3 as a thiol protecting agent analyzed via the DTNB assay

To analyze the efficiency of OPR3 to protect GSH in the presence of NEM, the thiol quantification method based on thiol reactivity of Ellman’s reagent (DTNB) was used with a standard curve generated with GSH. Both reagents were dissolved in DTNB assay buffer (120 mmol/L NaPi, pH 7.8, 6 mmol/L EDTA). The absorption change was analyzed by incubating 132 µL KPi buffer 40 mmol/L pH 8.0, 20 µL NADPH (2.4 mmol/L), 4 µL OPR3 (0.47 mg/ml) and 4 µL NEM (2.5, 5 or 10 mmol/L) for 20 min. To the assay 40 µL GSH (1 mmol/L) were added, and samples were incubated for another 10 min. The same procedure was also carried out without either OPR3 or NEM or without both NEM and OPR3 using corresponding buffer volumes to adjust concentrations. The samples were mixed with buffer and DTNB as described before for the determination of the thiol calibration curve with GSH standards (40 µL samples or standard, 140 µL DTNB buffer, 20 µL DTNB 6 mmol/L). The determination of free thiol content was based on absorbance read at 412 nm in a 96-well plate reader (Biotek). All incubations and measurements were performed at 25°C. Each sample was prepared four times and absorbance read for each in triplicate individually (n=12). The sample containing NADPH and GSH at a NEM/GSH ratio of 0 had an amount of 196.6±1.7 µmol/L free thiols which was set to 100%.

### Tryptophan fluorescence measurements

The quenching of OPR3 tryptophan fluorescence by binding of 12-OPDA and ASC was studied at 25°C using the SFM spectrofluorometer (SFM 25; Kontron Instruments) by adding 10 µL of 8.5 mmol/L 12-OPDA or ASC (dissolved in buffer, containing 10% EtOH) to 130 µL buffer (KPi 40 mmol/L; pH 7.2) containing 4.5 µg OPR3. After 2 min incubation, Trp fluorescence was excited at 284, and emission spectra were recorded between 300 and 400 nm using scan range of 100 nm and scan speed of 100 nm/min. The response was set to 8.0 and the high voltage to 500 with other settings being default. The control measurement was performed by adding 10 µL of buffer containing 10% EtOH to the OPR3 sample. Results were corrected for background fluorescence of ligands dissolved in buffer. The quenching curve of OPR3 by ascorbate was analyzed as follows. A quartz cuvette containing 100 µL buffer and 25 µL OPR3 (0.47 mg/ml) was supplemented with 10 µL of ASC (0 to 35 mmol/L in buffer). Immediately after mixing Trp fluorescence was recorded using settings as above. No pH change was observed upon ASC addition. Measurements were performed at least in triplicate.

### The interaction of 12-OPDA with ASC and GSH

The interaction of ascorbate with 12-OPDA was studied by means of mass spectrometry, isothermal titration calorimetry (ITC), and OPR3-coupled enzyme assay with glutathionylated 12-OPDA as control. The OPR3-coupled enzyme assay for detection of GSH- and ASC-modified 12-OPDA was performed spectrophotometrically (Shimadzu 2401) in a cuvette containing 15 µL NADPH, 0.5 µg OPR3 and 100 µL buffer. To start the NADPH-coupled reduction of un-modified 12-OPDA, 1 µL of 12-OPDA incubated with ASC or GSH was added. The ascorbate- and GSH-modified 12-OPDA samples were obtained by incubating 40 µL of ASC or GSH stocks (6.25 mmol/L, and 50 mmol/L in KPi 40 mmol/L pH 7.2) with 10 µL 12-OPDA (10 mmol/L) at 25°C on a shaker at 1500 rpm (Ika, VXR basic Vibrax). At indicated time intervals, samples were taken and placed on ice to stop the reaction. The same procedure was carried out with 12-OPDA dissolved in buffer as control. The final concentrations of 12-OPDA was 2 and 0 mmol/L (control), while it was 5 and 40 mmol/L for the redox buffers. Measurements were performed at least in triplicate and the rate for initial 30 seconds was considered for activity calculations. Activities were related to maximal activity obtained with 12-OPDA (control).

ITC analysis of the interaction between 12-OPDA and ASC was studied by injection of 5.88 mmol/L 12-OPDA into 0.318 mmol/L ASC. The compounds were dissolved in 0.1 M KPi pH 6.6. To ensure equal EtOH amounts in both syringe and cuvette, ascorbate was supplemented with solvent (EtOH 1% v/v). The total injection volume was 300 μL and the cell volume 1,400 μL. The parameters for each injection were set to 10 µL injection volume, 20 s duration, spacing between injections 60 s, 264 rpm stirring speed, cell temperature 25°C, reference power 15, feedback, high, fast equilibration auto. The respective heat changes recorded for the blank, i.e. injection of 5.88 mmol/L 12-OPDA into buffer were subtracted. The integrated heat responses were fitted to the single binding site model using the ORIGIN software package supplied with the calorimeter (MicroCal LLC ITC).

### Nano-UPLC/nano-ESI-MS analysis

For the MS analysis 40 µL of ASC (50 mmol/L in KPi, 40 mmol/L, pH 7.2) was mixed with 10 µL 12-OPDA (10 mmol/L) at 25°C for 18 h as described above and analysed via nano-HPLC/nano-ESI-MS. An Acquity UPLC M-Class® system (Waters Corp., Manchester, UK), consisting of µSample Manager, Trap Valve Manager, µBinary Solvent Manager und Auxiliary Solvent Manager was on-line coupled to a Synapt G2Si Q-IMS-*oa*-TOF mass spectrometer (Waters Corp.), equipped with a nano-ESI source. The trap column of the UPLC-system was a 2D Symmetry® C18, 5 µmol/L particles, 100 Å, 180 µmol/L x 20 mmol/L, the separation column a HSS T3, C18, 1.8 µmol/L particles, 75 µmol/L x 150 mmol/L (both Waters Corp.).

The 12-OPDA/ascorbate reaction product was diluted 1:100 using a solution of 3% acetonitrile in water, containing 0.1% trifluoroacetic acid. 2.5 µL of the mixture was injected into the UPLC-system. The sample was loaded onto the trap column for 2 min using 99.5% solvent A (water containing 0.1% formic acid) and 0.5% solvent B (acetonitrile, containing 0.1% formic acid) at a flow rate of 5 µL/min. The separation was performed at a flow rate of 300 nL/min, using a 60 min gradient (5 min 95% A, in 30 min to 65% A, in 5 min to 15% A, holding 15% A for 5 min, back to 95% A in 1 min, and 95% A for 14 min).

The mass spectrometer was operated in positive ion mode, TOF-mode, and resolution mode. Mass spectra were recorded in the range of *m/z* 350-2000. The nano-ESI Emitter was operated at 1.6 kV, sampling cone 25 V, source offset 55 V, source temperature 60°C. Lock mass spectra were recorded every 45 s for 0.5 s, using a mixture of 25 pg/µL Leucine Enkephaline [*m/z* 556,2766] and 200 fmol/µL GluFib in 50% acetonitrile/H_2_O with 0.1% formic acid, supplied by the Auxiliary Solvent Manager at a flow rate of 0.6 µL/min. The spray voltage of the lock mass sprayer was set to 3.2 kV. Data analysis and processing of the mass spectra was performed using MassLynx™ 4.1, (Waters Corp., Manchester, UK).

### Data processing, Databases and *in silico* studies

Data processing, calculations and presentations were performed using Microsoft Office 2010. Protein sequences, molar weight of OPR3 (42,691 g/mol) and PDB entries were obtained from Uniprot (http://www.uniprot.org). The superimposition of OPR3 (2g5w, green ribbon structure) onto that of rice MDHAR3 (herein referred as MDHAR, 5jcn, blue ribbon structure) was performed with integrated align command of PyMOL (The PyMOL Molecular Graphics System, Version 1.2r3pre, Schrödinger, LLC) using default settings.

### Plant growth and stress application

Arabidopsis (*Arabidopsis thaliana*) wild-type (Col-0) and *opr3* (Col-0) plants were grown for 3 and 6 weeks under day/night cycles of 10/14 h with light 100 µmol photons.m^-2^.s^-1^ at 22/19°C (day/night temperature) and 50% relative humidity. To enhance photorespiration, plants were transferred in an acrylic chamber continuously supplied with CO_2_-deprived air (sodalime, 9 h, 37 ppm CO_2_) in parallel, plants from the same set (WT and *opr3*) were placed in acrylic chambers continuously supplied with air (9 h, 460 ppm CO_2_). The CO_2_ content was determined using the BINOS 100 gas analyzer. After treatment rosettes were directly cut, frozen in nitrogen and stored at −80°C for 24-48 h. The ascorbate content was determined in the material that was pulverized in liquid nitrogen and then directly assayed (Kumar et al. 2019.

### Determination of photosynthetic parameters

Photosynthetic parameters of RES-treated leaves were analyzed as follows. Six leaf discs of 5 mmol/L diameter were cut out and placed immediately on 100 µmol/L CaCl_2_ solution (for membrane stabilization) containing 0.5 mmol/L RES compound (MVK and 12-OPDA, dissolved in EtOH). The control solution without RES contained EtOH instead of RES compounds. Petri dishes with floating leaf discs were closed with Petri dish lid and exposed to light (100 μmol photons m^−2^ s^−1^) at 22°C. The effective quantum yield of PS II (ΔF/Fm’) was measured using the photosynthesis yield analyzer (Mini-PAM, Walz, Germany) and calculated as (Fm’–F)/Fm’, where F is the fluorescence yield of the light-adapted sample in the steady state and Fm’ the maximum fluorescence yield after applying a saturating light pulse of 800 ms duration at an intensity of 3000 µmol photons.m^-2^.s^-1^ (Rascher et al. 2000).

## Supplementary Data

**Suppl. Fig. 1: ITC titration of 12-OPDA with ASC.**

**Suppl. Fig. 2: Nano-UPLC analysis of the 12-OPDA/ascorbate adduct**

**Suppl. Fig. 3: Mass spectrometric analysis of the 12-OPDA/ascorbate adduct**

**Suppl. Fig. 4: Mass spectrum of the** [**M+H**]**^+^-Ion of the 12-OPDA/ascorbate adduct**

**Suppl. Fig. 5: Reduction of the typical MDHAR substrate ferricyanide by OPR3**

**Suppl. Fig. 6: Oxygen electrode measurements of NADPH and CP in presence or absence of OPR3**

**Suppl. Table 1: Transcripts co-expressed with MDHAR3 and OPR3 from *A. thaliana***

## Supporting information

Maynard et al.-Supplementary Materials

## Acknowledgements

This work was supported by Deutsche Forschungsgemeinschaft (DI346/17&19). VK acknowledges funding from DAAD, Germany. We thank Dr. Stephan Unger, Faculty of Biology/Experimental Ecology, University of Bielefeld for provision of the BIOTEK CO_2_ gas analyzer.

