## Supplementary material for "12-oxophytodienoic acid reductase 3 (OPR3) functions as NADPH-dependent α,β-ketoalkene reductase in detoxification and monodehydroascorbate reductase in redox homeostasis": Maynard et al.-Supplementary Materials

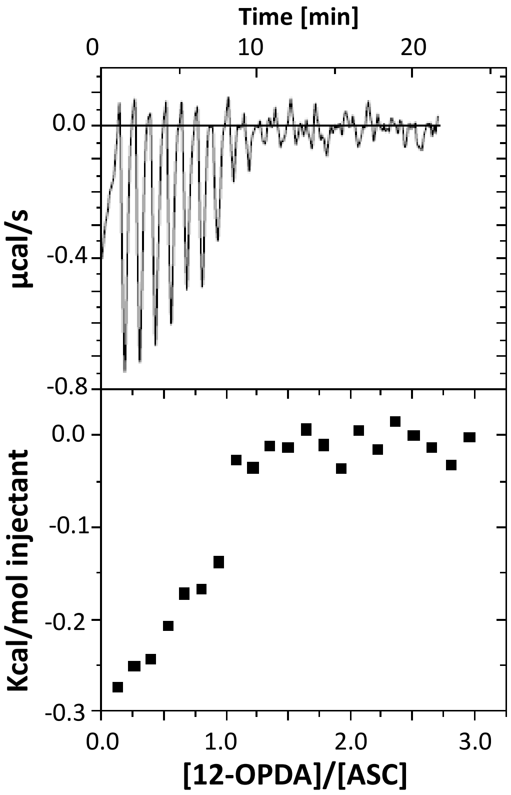


**Figure supplement S1. ITC analysis of 12-OPDA (10 µL of 5.88 mM) injected into ascorbate (318 µM, 1.4 mL).** Details see Materials and Methods

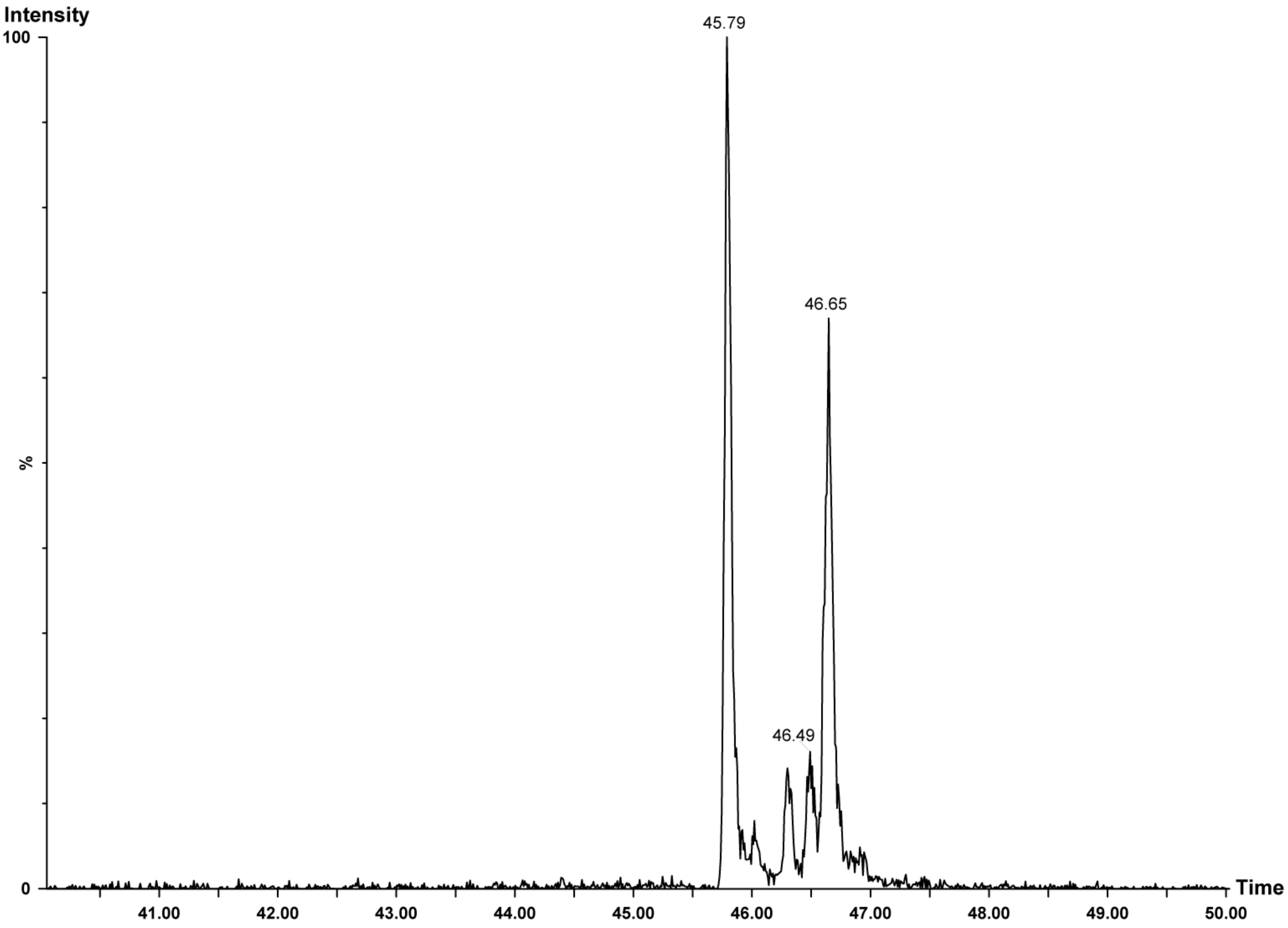


**Figure supplement S2. Chromatographic profile** of the nano-UPLC analysis of the 12-OPDA/ascorbate adduct

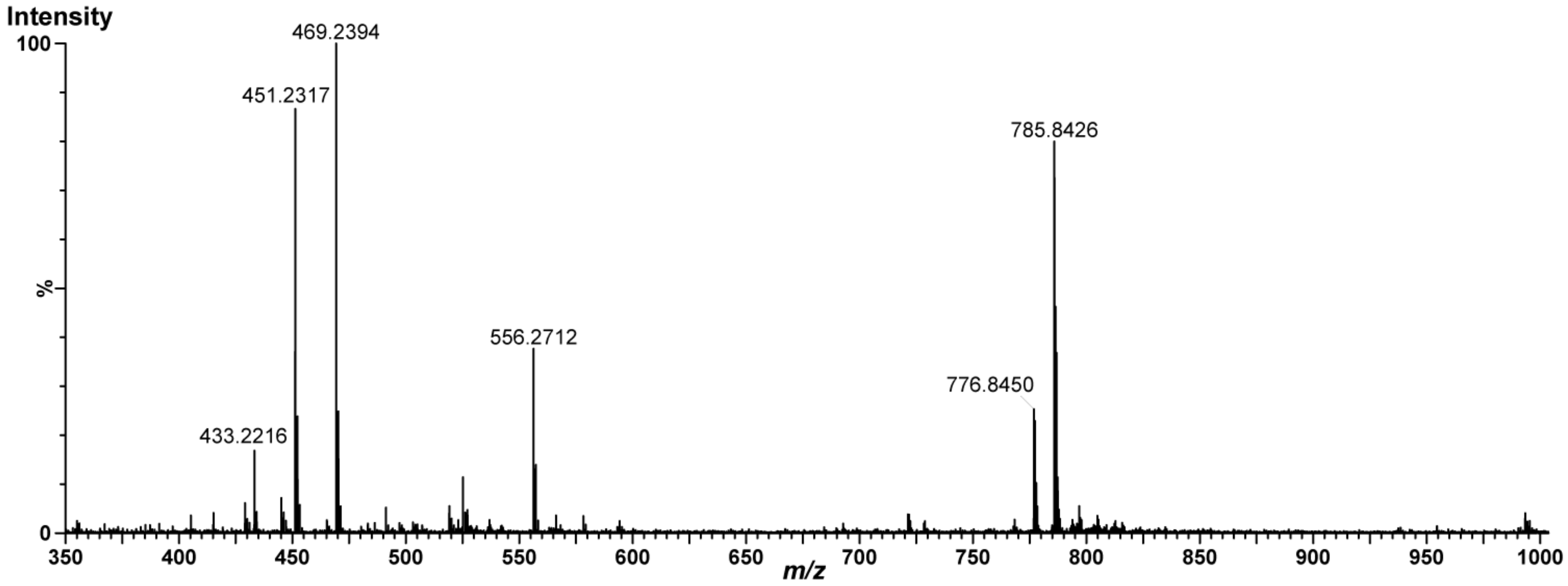


**Figure supplement S3. Mass spectrum of the chromatographic peak with a retention time of 45.79 min. [M+H]$^{+}$-ion of the covalent 12-OPDA/ascorbate adduct at *m/z* 469.2394**. The signal at *m/z* 451.2317 most likely corresponds to 12-OPDA/ascorbate adduct (minus 18 amu of lost water). Also detected were ions of the reference compounds (leucine enkephalin, [M+H]$^{+}$, *m/z* 556.2712; GluFib, [M+2H]$^{2+}$, *m/z* 785.8426) and a water loss of GluFib (*m/z* 776.8450).

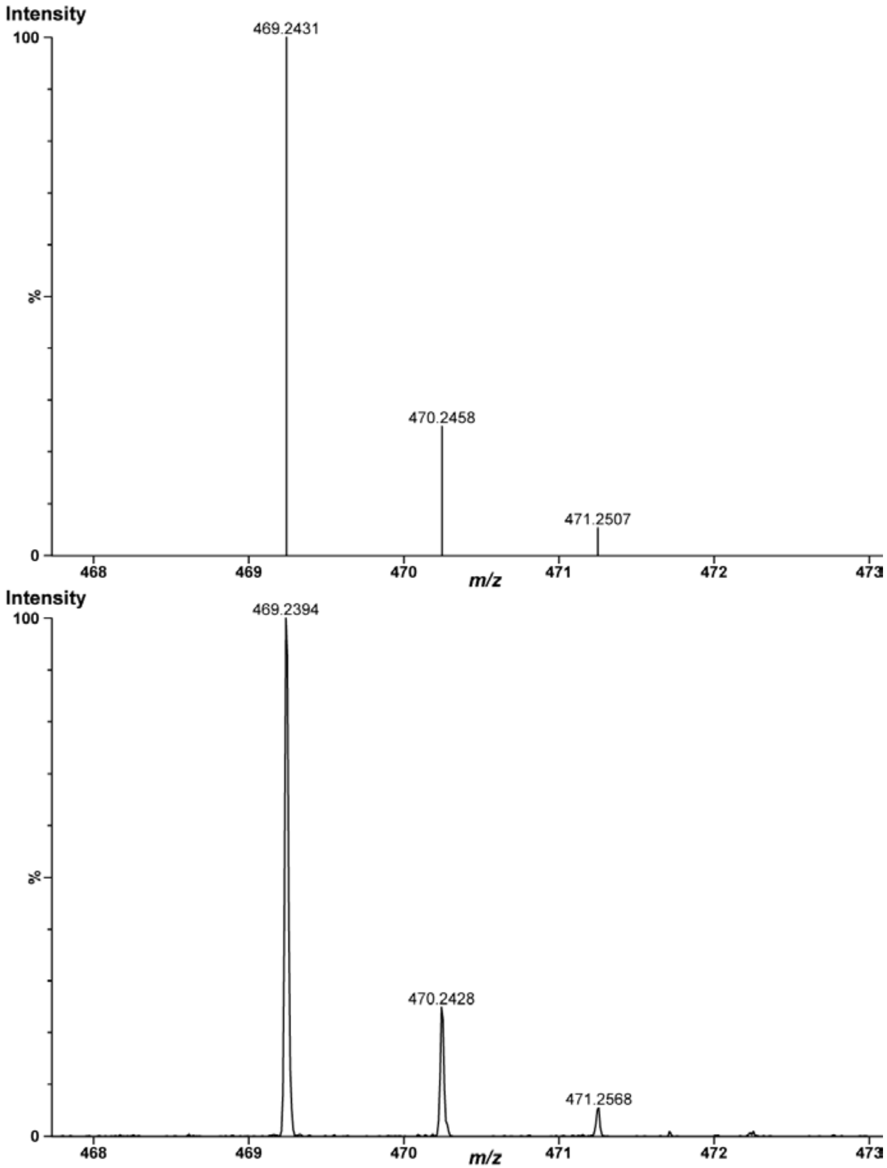


**Figure supplement S4. Mass spectrum and centroided mass spectrum (above) of the [M+H]$^{+}$-ion of the covalent 12-OPDA/ascorbate adduct** at *m/z* 469.2431 with its isotopic pattern.

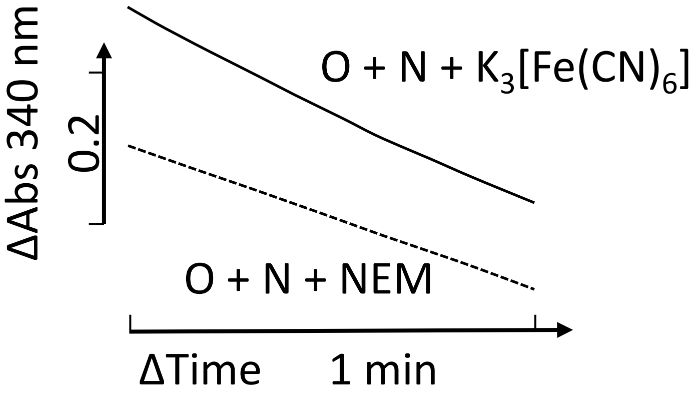


**Figure supplement S5. The typical MDHAR substrate $K_3[Fe(CN)_6]$ is effectively reduced by OPR3.** OPR3 (O) uses ferricyanide as substrate with similar rate as observed for NEM (570 µM in assay). No decrease in absorbance at 340 nm ocurred when NADPH (N) was absent and no OPR3 activity was observed when OPR3 enzyme assay was performed with ferrocyanide. OPR3 activity tests were performed as described in Materials and Methods section.

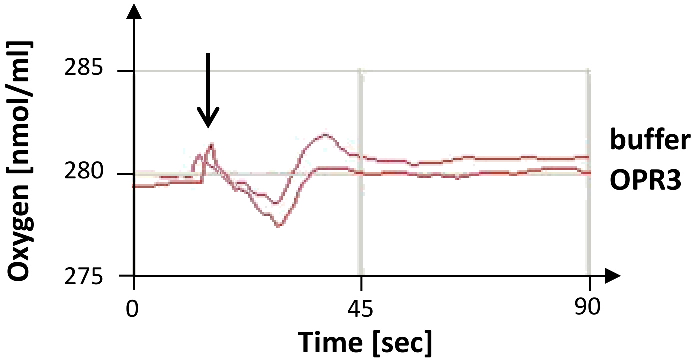


**Figure supplement S6. Oxygen recording of the assay containing NADPH and CP in the presence or absense of OPR3.** Overlaid sectors of oxygen electrode recordings of buffer (2.29 mL) containing NADPH (126 µM) and CP (742 µM), with 50µL buffer (upper recording) or OPR3 (0.2 µg/µL) added as indicated by arrow. The graphs are sections taken from 7 min recordings. Same results were obtained in multiple measurements. Details see Materials and Methods section

**Table supplement S1. Overlapping genes coexpressed with OPR3 (At2g06050) and MDHAR3 (At3g09940).** Transcriptomes were ranked by Mutual Rank (MR) and processed to reveal correlation by small rank values (Obayashi et al., 2017). Values taken from the ATTED-II website.

| genes coexpressed with OPR3 and MDHAR | function | MR, OPR3 as bait | MR, MDHAR as bait |
|---|---|---|---|
| At4g39030 | MATE efflux protein | 402 | 25.9 |
| At2g39420 | alpha/beta-Hydrolase protein | 22.5 | 77.5 |
| At1g23850 | | 749.9 | 16.2 |
| At3g25780 | allene oxide cyclase 3 | 29.3 | 42.7 |
| At5g19110 | | 468.2 | 24.1 |
| At1g76640 | Calcium-binding EF-hand protein | 274.2 | 72.8 |
| At3g51450 | Calcium-dependent phosphotriesterase protein | 3.9 | 279.7 |
| At3g21230 | 4-coumarate:CoA ligase 5 | 1116.9 | 323.5 |
| At2g29440 | glutathione S-transferase tau 6 | 843.4 | 686 |
| At2g30870 | glutathione S-transferase PHI 10 | 708.3 | 75.9 |
| At5g60300 | Concanavalin A-like lectin protein kinase | 2270.9 | 647.3 |
| At5g05600 | 2-oxoglutarate (2OG) and Fe(II)-dependent oxygenase | 15.9 | 614 |
| At3g44320 | nitrilase 3 | 1692.5 | 558.4 |
| At2g27690 | | 41.2 | 199.5 |
| At4g27860 | vacuolar iron transporter (VIT) | 1658.5 | 118.3 |
| At3g50280 | HXXXD-type acyl-transferase | 219.7 | 71.3 |
| At1g28480 | Thioredoxin | 78.5 | 185.9 |
| At5g09980 | elicitor peptide 4 precursor | 1828.2 | 580.3 |
| At5g05730 | anthranilate synthase alpha subunit 1 | 224.8 | 361.5 |
| At3g23250 | myb domain protein 15 | 224.8 | 361 |
| At5g67080 | mitogen-activated protein kinase kinase kinase 19 | 449.8 | 486 |
| At5g13220 | jasmonate-zim-domain protein 10 | 8.0 | 626 |
| At1g19180 | jasmonate-zim-domain protein 1 | 48.4 | 505 |
| At3g19010 | 2-oxoglutarate (2OG) and Fe(II)-dependent oxygenase | 289.6 | 366.2 |
| At4g39950 | cytochrome P450. family 79. subfamily B. polypeptide 2 | 660.3 | 441.7 |
| At3g54640 | tryptophan synthase alpha chain | 2143.5 | 164.2 |
| At1g20510 | OPC-8:0 CoA ligase1 | 1.1 | 464.4 |
| At5g07460 | peptidemethionine sulfoxide reductase 2 | 588 | 314.4 |
| At2g35930 | plant U-box 23 | 690.1 | 207.9 |
| At1g61120 | terpene synthase 04 | 37.4 | 826.6 |
| At4g10390 | Protein kinase | 394 | 188.7 |
| At4g31500 | cytochrome P450. family 83. subfamily B. polypeptide 1 | 209.9 | 617.5 |
| At1g74950 | TIFY domain/Divergent CCT motif | 10.2 | 814.8 |
| At1g74100 | sulfotransferase 16 | 40.1 | 778.1 |
| At1g10700 | phosphoribosyl pyrophosphate (PRPP) synthase 3 | 1133.9 | 247.2 |
| At4g30530 | Class I glutamine amidotransferase-like | 21.3 | 747.6 |
| At4g12720 | MutT/nudix protein | 286 | 382.5 |
| At2g42760 | | 43.9 | 579.4 |
| At5g45280 | Pectinacetylesterase | 1060.6 | 474.6 |
| At1g67560 | PLAT/LH2 domain-containing lipoxygenase | 1348.8 | 2222.7 |
| At1g30135 | jasmonate-zim-domain protein 8 | 23 | 1056.4 |
| At1g77420 | alpha/beta-Hydrolase | 382.2 | 1215.1 |
| At3g49620 | 2-oxoglutarate (2OG) and Fe(II)-dependent oxygenase | 1039.3 | 244.6 |
| At3g09830 | Protein kinase | 226.7 | 621.2 |
| At5g36880 | acetyl-CoA synthetase | 3196.3 | 204.3 |
| At1g17750 | PEP1 receptor 2 | 83.6 | 313.2 |
| At5g49280 | hydroxyproline-rich glycoprotein | 1929.1 | 1877.4 |
| At2g38240 | 2-oxoglutarate (2OG) and Fe(II)-dependent oxygenase | 106.5 | 1201.5 |
| At1g59870 | ABC-2 and Plant PDR ABC-type transporter | 840.8 | 1220.8 |
| At3g01830 | Calcium-binding EF-hand protein | 231.4 | 707.3 |
| At5g22630 | arogenate dehydratase 5 | 130 | 929.4 |
| At3g50930 | cytochrome BC1 synthesis | 175.9 | 513.4 |
| At4g18950 | Integrin-linked protein kinase | 408.7 | 364.1 |
| At2g32140 | transmembrane receptor | 150.9 | 754.4 |
| At1g06620 | 2-oxoglutarate (2OG) and Fe(II)-dependent oxygenase | 43.7 | 11.9 |
| At5g36220 | cytochrome p450 81d1 | 102.9 | 1310.8 |
| At5g38710 | Methylenetetrahydrofolate reductase | 246.6 | 327.3 |
| At5g11670 | NADP-malic enzyme 2 | 1474.9 | 571.8 |
| At5g03630 | Pyridine nucleotide-disulphide oxidoreductase | 288.1 | 141.3 |
| At1g76040 | calcium-dependent protein kinase 29 | 938.2 | 520.8 |
| At2g24850 | tyrosine aminotransferase 3 | 136.1 | 36.6 |
| At1g26730 | EXS (ERD1/XPR1/SYG1) | 153.3 | 1441.8 |
